# FUNCTIONAL ANALYSIS OF BIPARTITE NRF2 ACTIVATORS THAT OVERCOME FEEDBACK REGULATION FOR AGE-RELATED CHRONIC DISEASES

**DOI:** 10.1101/2025.05.30.656434

**Authors:** Dmitry M. Hushpulian, Navneet Ammal Kaidery, Priyanka Soni, Andrey A. Poloznikov, Arpenik A. Zakhariants, Alexandra V. Razumovskaya, Maria O. Silkina, Vladimir I. Tishkov, Eliot H. Kazakov, Abraham M. Brown, Irina N. Gaisina, Young-Hoon Ahn, Sergey V. Kazakov, Nancy Krucher, Sudarshana M. Sharma, Bindu D. Paul, Irina G. Gazaryan, Sergey V. Nikulin, Bobby Thomas

## Abstract

Activating Nrf2 with small molecules is a promising strategy for countering aging, oxidative stress, inflammation, and various disorders, including neurodegeneration. The primary regulator of Nrf2 protein stability is Keap1, a redox sensor protein and an adapter in the Cullin III ubiquitin ligase complex, which labels Nrf2 for proteasomal degradation. The known Nrf2 activators either chemically modify sensor thiols in Keap1 or competitively displace Nrf2 from the ubiquitin ligase complex. The latter approach is considered the most suitable for continuous administration, as non-specific chemical modifiers of Keap1 thiols also modify active thiols on other proteins, thus causing side effects. However, when transitioning from homogeneous to cell-based assays, genuine displacement activators show a significant loss in potency by several orders of magnitude. As we demonstrate here, this offset is due to the presence of high micromolar concentrations of Keap1 in both the cell lines and brain tissue. A potential solution could involve targeted delivery of an alkylating agent to Keap1 to achieve the desired specificity. Transcriptomic analysis of a cell-permeable Nrf2 peptide bearing an alkylating fumarate moiety indicates selective activation of the Nrf2 genetic program, confirming the high specificity of this approach. The Nrf2-triggered genetic program has a feedback regulation mechanism through the activation of Bach1, an Nrf2 transcriptional repressor, which is elevated in age-related neurodegeneration. Thus, a benign bipartite Nrf2 activator with Bach1 inhibition properties is needed for maximal benefits. The recently developed heterocyclic carboxamide, HPPE, shows overlap with the Nrf2 pathway activated by the fumarate-linked Nrf2 peptide and with zinc and tin protoporphyrins, which are recognized inhibitors of Bach1. Therefore, HPPE presents a promising and unique combination of the two desired activities that could be further optimized to treat age-related neurodegeneration.

**Highlights:**

- The decrease in potency for reversible displacement activators of Nrf2 in biological assays is attributed to high micromolar concentrations of Keap1 and competition with endogenous Keap1 client proteins.
- Nrf2 activators specific for Keap1 should combine a displacement scaffold with a substitution that undergoes intracellular conversion into active pro-oxidant or alkylating species.
- Cell-permeable fumarate-linked Nrf2 peptide solely activates the Nrf2 antioxidant genetic program, as demonstrated by transcriptomic analysis.
- HPPE, a small bipartite molecule, exhibits properties of both Nrf2 activation and Bach1 inhibition, to bypass feedback regulation by targeting both Keap1 and Bach1.

## 1. Introduction

The trend of a growing global aging population is accompanied by the development of numerous health concerns, with aging-related diseases being most prominent. As aging is the most significant risk factor for chronic neurodegenerative diseases, a comprehensive understanding of aging mechanisms and their link to related diseases is crucial for developing effective preventive and therapeutic strategies (Hou et al. 2019). Aging is a natural physiological process resulting in a gradual decline in the body’s physiological functions. This process is based on complex molecular and cellular changes, including mitochondrial dysfunction, DNA damage, telomere shortening (referred to as the Hayflick limit), lipid peroxidation, and protein oxidative modifications. A fundamental theory of aging involves the free radical theory, first introduced by Denham Harman (Harman 1956), who proposed that aging results from accumulated damage caused by free radicals generated by metabolic reactions or by-products of cellular processes. Later, Harman highlighted the significant role of mitochondrial dysfunction in aging (Harman 1972). The mitochondrial theory of aging has been debated; however, mitochondrial dysfunction is a consistent and conserved primary hallmark of aging (Santos, Sinha, and Lindner 2018) (Bartman, Coppotelli, and Ross 2024). Mitochondrial respiration is inseparable from its by-products, reactive oxygen species (ROS), which leak from Complex I and Complex III. As such, it must be supported by an effective antioxidant system that detoxifies ROS. Moderate ROS levels modulate metabolism and activate the antioxidant genetic program, whereas acute ROS overloads the cells’ redox capacity, leading to cellular dysfunction and death (Santos, Sinha, and Lindner 2018), (Davalli et al. 2016). Accumulating oxidative damage to cellular components is known to interfere with cell division, leading to cancer, apoptosis, or cellular senescence, which accumulates with age (Davalli et al. 2016) (Yang et al. 2024). The brain is especially vulnerable to oxidative damage because of its lipid-rich nature and high metabolic activity that results in increased production of ROS. Unsurprisingly, age-related neurodegenerative diseases are directly linked to mitochondrial health, cellular capability to detoxify ROS, and the potential to clear protein aggregates in neural cell lineages.

Cellular redox homeostasis is regulated by the intrinsic antioxidant genetic program, orchestrated by the key transcription factor nuclear factor erythroid 2-related factor 2 (Nrf2). Nrf2 is constitutively synthesized in the cell and, in the absence of oxidative or xenobiotic stress, undergoes proteasomal degradation through its interaction with the adapter protein Keap1, a redox sensor in the ubiquitin ligase Cul III complex. Two other stress-induced mechanisms for fine-tuning Nrf2 protein stability occur via beta-transducin repeat-containing E3 ubiquitin-protein ligase (β-TrCP) and E3 ubiquitin-protein ligase synoviolin (Hrd1), which play a secondary role (Harder et al. 2015). Keap1 has a set of highly reactive thiols, which, when modified, disrupt the Nrf2/Keap1 or Keap1/Cul III interactions, resulting in the stabilization of the Nrf2 protein. The Nrf2-mediated genetic program has inhibitory feedback regulation by upregulating the expression of Keap1 (Lee et al. 2007) and the transcriptional repressor Bach1 (Jyrkkanen et al. 2011). Maintaining a proper Keap1/Nrf2 balance is crucial for autophagy and aggregate clearance (Kopacz et al. 2020). The Nrf2 activation mechanism through Keap1 binding affects the processing of another adaptor protein, p62 (also known as SQSTM1), as the latter is both a client of Keap1 and a target gene of Nrf2. A regulatory loop is formed between these two essential signaling pathways: the Nrf2-triggered genetic program and p62-supported mitophagy (Gureev et al. 2020). A recent comprehensive dynamic model of ROS action network relevant to Parkinson’s disease onset with aging (Kolodkin et al. 2020) took into account a sophisticated defense network including the Keap1/Nrf2 axis, p62 in mitophagy, nuclear factor kappa-light-chain-enhancer of activated B cells (NF-κB) modulation, and the stress sensor DJ1, and identified the targets for “life-extending interventions”: mitochondrial synthesis, Keap1 degradation, and p62 metabolism. In other words, Nrf2 activation was again pointed to as a potential magic bullet for increasing lifespan and counteracting neurodegeneration. Nrf2 is also known to play a key role in reversing nuclear aging defects in premature aging, as well as in other age-related diseases (Kubben et al. 2016). Therefore, Nrf2 is a well-justified target for medical intervention to prevent and treat age-related diseases of various etiologies.

Multiple approaches have been followed to harness the Nrf2 pathway for therapeutic benefits. Activation of the Nrf2-driven genetic program with small molecules that modify thiols on Keap1 is a popular approach, despite the existence of unavoidable side effects originating from the non-specific thiol modification by common Nrf2 activators, such as curcumin (turmeric) and sulforaphane (broccoli), or by FDA-approved medications based on fumarate esters (Tecfidera, Vumerity, Bafiertam) (Dinkova-Kostova and Copple 2023). The requirements for Nrf2 activators intended to treat ongoing neurodegeneration must be stringent, as non-specific thiol modifications can result in non-specific oxidation of cellular components and increased ROS generation, which are already associated with ongoing neurodegeneration. A benign solution to the problem could be the development of competitive and reversible Nrf2 displacement activators. However, the current generation of displacement activators with high Keap1 binding affinity fails to display low EC_50_ values in cell-based assays; therefore, high doses of displacement activators are required for *in vivo* testing. To gain a precise understanding of the problem of low effectiveness of Nrf2 displacement activators, we employed a range of various novel small molecules and biologics, experimental methodologies, including a proprietary reporter assay, protein quantification, and transcriptomic approaches, to uncover the roots of the existing problem and formulate the requirements for developing an ideal Nrf2 activator. Our findings lay the groundwork for developing new therapeutics that target the Nrf2 signaling cascade in aging and neurodegenerative diseases, while minimizing side effects and enhancing therapeutic benefits.

## 2. Materials and Methods

### 2.1. Animals

We used male and female C57BL/6J mice (Jackson Laboratories, https://www.jax.org/strain/000664) in the present study. Mice were housed in a pathogen-free facility in humidity and temperature-controlled rooms maintained under a 12-hour light/dark cycle. They were given free access to standard rodent chow and water. All experimental procedures were conducted according to NIH Guidelines for the Care and Use of Laboratory Animals. The Institutional Animal Care and Use Committee of the Medical University of South Carolina approved all the procedures.

### 2.2. Peptides, Nrf2 activators and Bach1 inhibitors

*N*,*N*’-(Naphthalene-1,4-diyl)bis(4-methoxybenzsulfonamide) (NMBSA) was purchased from Sigma-Aldrich (USA), 4-methyl-*N*-(quinolin-8-yl)benzenesulfonamide (8TQ), *N*-(quinolin-8-yl)-2,4,6-trimethylbenzenesulfonamide (8QBSA), *N*-(5-bromoquinolin-8-yl)-2,4,6-trimethylbenzenesulfonamide (Br-8QBSA), *N*-(5-chloroquinolin-8-yl)-2,4,6-trimethylbenzenesulfonamide (Cl-8QBSA) were purchased from ChemDiv (USA). CPUY192018, zinc protoporphyrin (ZnPP) were purchased from MedChemExpress (USA), tin protoporphyrin (SnPP) - from Cayman Chemical (USA). All other reagents were from Sigma-Aldrich (St Louis, MO). *N*-(2-(2-Hydroxyethoxy)ethyl)-1-methyl-2-((6-(trifluoromethyl)benzo[*d*]thiazol-2-yl)amino)-1*H*-benzo[*d*]imidazole-5-carboxamide (HPPE) (purity 99.8%) was custom synthesized by Pharmaron (USA). Cell-permeable variants of Nrf2 peptides (>99% purity) – YGRKKRRQRRRAQLQLD**EETGE**FLPIQ (wild-type), YGRKKRRQRRRAQLQLD**PETGE**FLPIQ (mutant), and YGRKKRRQRRRAQLQLD**PETGE**FLPIQK-NHCO-fumarate methyl (fumarate-linked peptide) were custom synthesized by LifeTein LLC (NJ, USA) with >95% purity. The peptides were tested in a fluorescence polarization (FP) assay using recombinant Keap1 to ensure that no decrease in affinity for the Kelch domain occurred upon the addition of a TAT sequence to the ETGE motif peptides used in this work. The competitive displacement activity of cell-permeable peptides was evaluated using the Keap1-Nrf2 inhibitor screening assay kit (BPS Biosciences, San Diego, CA) according to the manufacturer’s instructions, as previously described by us (Ahuja et al. 2021), which yielded dissociation constant (K_D_) values within the 50-80 nM range.

### 2.3. Neh2-luc reporter assay

SH-SY5Y cell line stably expressing Neh2-luc (Smirnova et al. 2011) reporter was grown in DMEM/F12 supplemented with GlutaMAX (Thermo Fisher Scientific) containing 10% FBS, 100 U/mL penicillin, and 100 μg/mL streptomycin) and plated into 96-well white flat-bottom plates at 15,000 cells/well in 100 μL serum and incubated for 16 h at 37 °C, 5% CO2. The small-molecule compounds were prepared as 10 mM stock solutions in dimethyl sulfoxide (DMSO). Then, the compounds were added to the wells, resulting in final concentrations ranging from 2.5 to 40 μM. Stock solutions of the peptides were prepared in 5 mM Tris-HCl buffer, pH 7.0, at concentrations of 4 or 8 mM, and were added to the wells to achieve the desired final concentration. The plates were incubated for three hours at 37 °C. The medium was removed, and cells were lysed with 20 μL of Lysis buffer (Promega) for 7 min at room temperature (RT). Luciferase activity was then measured using a GloMax multidetection plate reader (Promega) with 80 μL of BrightGlo™ reagent (Promega, Madison, WI). The reporter activation was normalized to the background (where only DMSO was added). Tert-butylhydroquinone (TBHQ) was used as a positive control. For the time-course experiments, two μL aliquots of the compounds or 5 μL aliquots of peptides were added to the wells at different time points, and the assay was conducted as described above. The experiments were performed in triplicate.

### 2.4. RT-PCR for Nrf2 target genes

Wild-type (WT) or Nrf2 knockout (KO) mouse embryonic fibroblast (MEF) was treated with wild-type TAT-Nrf2-peptide (200 µM) or tert-butyl hydroquinone (TBHQ, 20 µM) for 4 or 16h. Total RNA was isolated using TRIzol reagent (Invitrogen) according to the manufacturer’s protocol. One µg of total RNA was reverse transcribed using a High-Capacity cDNA Reverse Transcription Kit (Invitrogen). cDNA was diluted, and 20 ng was used to amplify in an ABI prism 7900HT Real-time PCR system (Applied Biosystems) for transcripts of Nrf2 dependent genes: Heme Oxygenase 1 (HMOX1, 5’-GGGTGATAGAAGAGGCCAAGA-3’ and 5’-AGCTCCTGCAACTCCTCAAA-3’) and NAD(P)H quinone oxidoreductase 1 (NQO1, 5’-AGCGTTCGGTATTACGATCC-3’ and 5’-AGTACAATCAGGGCTCTTCTCG-3’) using Fast SYBR® Green Master Mix (Invitrogen). Cycling parameters were 95°C for 10 s, followed by 60°C for 1 min for 40 cycles. Relative expression was calculated using the ΔΔCt method (Schmittgen and Livak 2008). Values are expressed as a fold of control reaction and normalized to beta-actin (Actb, 5’-CTAAGGCCAACCGTGAAAAG-3’ and 5’-ACCAGAGGCATACAGGGACA-3’) expression.

### 2.5. Keap1 and Nrf2 protein quantification assay

Since human and mouse Keap1 and Nrf2 have 94% and 99% homology, respectively, human recombinant Keap1 and Nrf2 proteins were used as standards and were produced as follows. The plasmids pET28a-His6-Halo-Tev-Keap1 and pET28a-His6-Halo-TEV-Nrf2 were purchased from Addgene (Parvez et al. 2015) and expressed in BL21(DE3) *E. coli cells* (C2527I, New England BioLabs), using 25 μM IPTG for induction. The bacterial pellet was collected and lysed in 50 mM Tris buffer, pH 8, containing 100 mM NaCl, 0.2 mg/ml lysozyme, and PMSF. His-tagged proteins were then isolated using the HisPur Ni-NTA purification kit (#88229, Thermo Scientific) with 200 mM Imidazole as the eluent in the same buffer. The eluted proteins were desalted using PD10 columns (#17-0851-01, GE Healthcare) and concentrated with Vivaspin6 30kDa MWCO spin cartridges (28-9323-17, GE Healthcare). The protein concentrations of Keap1 and Nrf2 were determined using Nanodrop, taking into account the extinction coefficients of 13.8 and 8.58 at 280 nm, and the molecular weights of 106.6 and 104.72 kDa (Lau et al. 2013), respectively. Samples for western blotting were prepared from the Neh2-luc reporter line and plain SH-SY5Y cells grown in DMEM/F12 medium supplemented with 10% fetal bovine serum (FBS) and antibiotics. For Neh2-luc, the media was supplemented with 500 μg/ml geneticin (G418, Sigma-Aldrich). Cell numbers were estimated with a TC10 cell counter (BioRad), and 100,000 cells were loaded per lane in a 12% TGX gel. Tissue lysates were prepared in TNES (Tris 50 mM, NaCl 150 mM, EDTA 5 mM, SDS 1%, NP-40 0.5%, Na-deoxycholate 0.5%, pH 7.4) buffer containing 1X protease inhibitor (P8340, Sigma), and the concentration was determined using the BCA assay. Brains (cortex, brainstem, and spinal cord) were collected from young (3-month-old) or old (15-month-old) C57BL/6J mice. In 12% TGX gels (Criterion, BioRad), 100 μg of brain lysates was separated along with standard proteins. Either 250, 100, 25, 10, 1, femtomoles of purified proteins were used as standards for Nrf2 and Keap1, respectively. After transferring to a nitrocellulose membrane, blots were probed with antibodies recognizing both human and mouse Keap1 (A21724, Abclonal) and human and mouse Nrf2 (A0674, Abclonal), and detected using appropriate HRP-conjugated secondary antibodies. Band intensity was measured using ImageLab software (Bio-Rad).

### 2.6. RNA-seq and Bioinformatics

Neuroblastoma SH-SY5Y cells were grown in 6-well plates in 2 mL medium per well up to 80% confluence. Cells were treated with either 100 μM of the peptide or 5 μM HPPE, zinc, or tin protoporphyrin, and DMSO was used as a control. After five hours of incubation, the medium was removed, and the cells were washed three times with DPBS. They were then lysed in 700 μL QiaZol. RNA isolation was performed using the miRNeasy Micro Kit (Qiagen, Germany) according to the manufacturer’s protocol. Nanodrop ND-1000 (Thermo Fisher Scientific, USA) was used to assess the quantity and purity of the extracted RNA. The quality control (QC) for the RNA was performed using an Agilent 2100 Bioanalyzer (Agilent Technologies, USA).

The libraries for mRNA sequencing were prepared from total RNA samples using the Illumina Stranded mRNA Library Prep Kit (Illumina, USA). Each sample was sequenced on the NextSeq 550 (Illumina, USA) to generate single-end 75-nucleotide reads. In the experiment with the small molecule compounds, the libraries for mRNA sequencing were prepared from total RNA samples using the MGIEasy RNA Library Prep Set (MGI Tech Co., China). The sequencing was conducted on the DNBSEQ-G400 (MGI Tech Co., China) to generate single-end 100-nucleotide reads.

The quality of FASTQ files was assessed with FastQC v0.11.9 (Babraham Bioinformatics, UK) and multiQC v1.9 (Ewels et al. 2016). The adapters were trimmed using FastP 0.21.1 (PMID: 30423086 (Chen et al. 2018)). The trimmed mRNA-seq reads were mapped to the reference human genome GENCODE release 37 (GENCODE GRCh38.primary assembly) using STAR 2.7.7a (Dobin et al. 2013). GENCODE release 37 genome annotation (gencode.v37.primary.assembly.annotation) (Frankish et al. 2019) was used to generate the count matrix with the featureCount tool from subread-2.0.1 aligner (Liao, Smyth, and Shi 2013), (Liao, Smyth, and Shi 2014).

The combined raw counts from RNA-sequencing data were processed for batch correction using ComBat (Zhang, Parmigiani, and Johnson 2020). Elbow-plot was used to determine the number of clusters for K-means clustering. Heatmaps were generated using the Interactive Complex Heatmap package (Gu and Hubschmann 2022). We used Enrichr (Chen et al. 2013) to evaluate pathways affected in each cluster broadly. For WGCNA analysis, the RNA-sequencing data were normalized using the Variance Stabilizing Transformation (VST) of the DESeq2 package, and the 75th quantile normalized data were used to construct the networks (Langfelder and Horvath 2008). The network was divided into modules based on correlation coefficient information at a soft power threshold of 10. We used dynamic tree cut to determine discrete modules containing genes with similar expression patterns, which were assigned an ID and color. After defining discrete modules using the dendrogram, we utilized the module Eigengene to correlate with the sample types and treatments (targets: Nrf2 or Bach1; treatments: Nrf2-peptide, HPPE, zinc-, and tin-protoporphyrin).

Differential expression analysis was conducted using DESeq2 v1.44.0 (Love et al., 2014). False discovery rates (FDRs) were calculated using the Benjamini–Hochberg procedure. To assess the statistical significance of differences in gene expression, FDR p-values with a threshold level of 0.05 were used. Gene Ontology enrichment analysis (Ashburner et al. 2000) (Gene Ontology et al. 2023) was performed with a built-in analysis tool on the GO web page (Thomas et al. 2022). The RNA-sequencing data generated in this study are available in the Gene Expression Omnibus (GEO) repository (GSE271364 and GSE287793).

### 2.7. Computer modeling

Small-molecule docking experiments were performed using the CDOCKER algorithm, as implemented in the Discovery Studio software suite (BIOVIA, San Diego, CA), followed by force field minimization and calculation of binding energies. The Nrf2 crystal structure with the bound inhibitor (4IQK.pdb) served as the starting template for this study. All ligands were imported and minimized using the ‘Prepare ligands’ protocol after adding hydrogen bonds. Force field minimization was performed using the molecular mechanics algorithm CHARMM, as implemented in Discovery Studio. A series of 50 positions was analyzed for each compound. The percentage of high-interaction energy positions that coincide with those of NMBSA in the crystal structure has been calculated and used as a comparative measure of compound affinity for the Kelch domain. Peptide docking experiments were performed using Discovery Studio, which employed the build protein and superimposition tools (versus the initial peptide in the crystal structure 5WFV.pdb), followed by manual attachment of fumarate to the peptide chain. Final peptide pose selection was performed by applying CHARMM minimization algorithms. The Peptide docking benchmark was created with a set of Scoring and Analysis protocols. Docking energies for the versions of cell-permeable Nrf2 TAT-peptides studied in the work, in comparison to the control ETGE peptide, showed no principal change (Table S1).

### 2.8. Statistical Analysis

Statistical analyses and graphs were performed using GraphPad Prism version 10.0.0 for Mac OS (GraphPad Software, San Diego, California USA, www.graphpad.com). One-way or two-way ANOVA with appropriate post hoc analysis was used for multiple comparisons, whereas an unpaired, two-tailed Student’s t-test was used for comparing two groups. All data were plotted as mean ±SEM and were considered significant when P ≤ 0.05.

## 3. RESULTS AND DISCUSSION

### 3.1. Low effectiveness of true displacement activators in the cell-based assay

There are two fundamentally different cell-based screening assays for Nrf2 activators. One approach is to use a transcription reporter, where a reporter gene is cloned under a promoter activated by Nrf2, as in the antioxidant response element (ARE)-luc reporter. The second method utilizes a fusion reporter with constitutive expression of a fusion protein containing the Neh2 domain of the Nrf2 protein, the first N-terminal domain, which serves as a recognition domain for Kelch domains in Keap1 dimer (see Nrf2 and Keap1 domain structures in Fig.1 A and B, respectively) in frame with firefly luciferase. The Neh2-luc fusion protein expressed undergoes the same fate as endogenous Nrf2, i.e., the fusion protein, recognized by Keap1, undergoes ubiquitination and proteasomal degradation (Fig.1C). The advantage of the fusion reporter assay is its immediate response to the inhibitors of Nrf2-Keap1 interaction and the possibility of real-time monitoring of reporter activation (Smirnova et al. 2011). In contrast, ARE-luc reporters require a significantly longer time to demonstrate activation, and the amplitude of the observed activation effect is much lower (see the comparison in Fig.1B in (Smirnova et al. 2011). Typical irreversible Nrf2 activators, which function by chemically modifying key Keap1 thiols, exhibit more than a 20-fold activation effect within three hours of incubation with Neh2-luc reporter cells, with EC_50_ values reflecting the potency of alkylating agents. As shown in Fig. 2A, bardoxolone and auranofin are among the most potent alkylating Nrf2 activators, exhibiting activity in the nanomolar range and becoming toxic above 1 μM. In contrast, dimethyl fumarate (DMF) and *tert-*butylhydroquinone (TBHQ) are much “softer” and become toxic at concentrations above 40 μM (Fig. 2B). Irreversible modification in the case of non-specific alkylating agents proceeds on a stoichiometric basis and targets all active thiols in all the proteins, including Keap1. Hence, this is a reason for the unavoidable toxicity of Nrf2-activating alkylators. However, “benign” displacement activators, such as those shown in Fig. 2C, demonstrate orders of magnitude lower potency in the Neh2-luc assay than in the homogeneous fluorescence polarization assay (FP assay). This is in accordance with the published literature, where NMBSA and CPUY have K_Ds of_ approximately 1 μM and 40 nM, respectively. However, EC_50_ in the Neh2-luc assay is above 30 μM for both compounds. The observed plateau (Fig. 2D), which never reaches the maximum activation threshold observed for irreversible activators (Fig. 2A and B), suggests that the system re-equilibrates for competitive displacement activators. The observed offset of EC_50_ by orders of magnitude for displacement activators (Fig. 2C) aligns with the published data on their biological activity in the ARE-luc assay and RT-PCR (Lu et al. 2016). The offset of the biologically active concentrations with respect to K_D_ values determined in the FP assay is typical for all displacement activators (see (Plusa 2022) references therein), which originates from competition under cellular conditions. In other words, the cellular abundance of Keap1 far exceeds the K_D_ values determined in FP assays, and, in addition, one may expect the abundance of other Keap1 client proteins, competing with the displacement activators besides Nrf2.

**Fig. 1.**
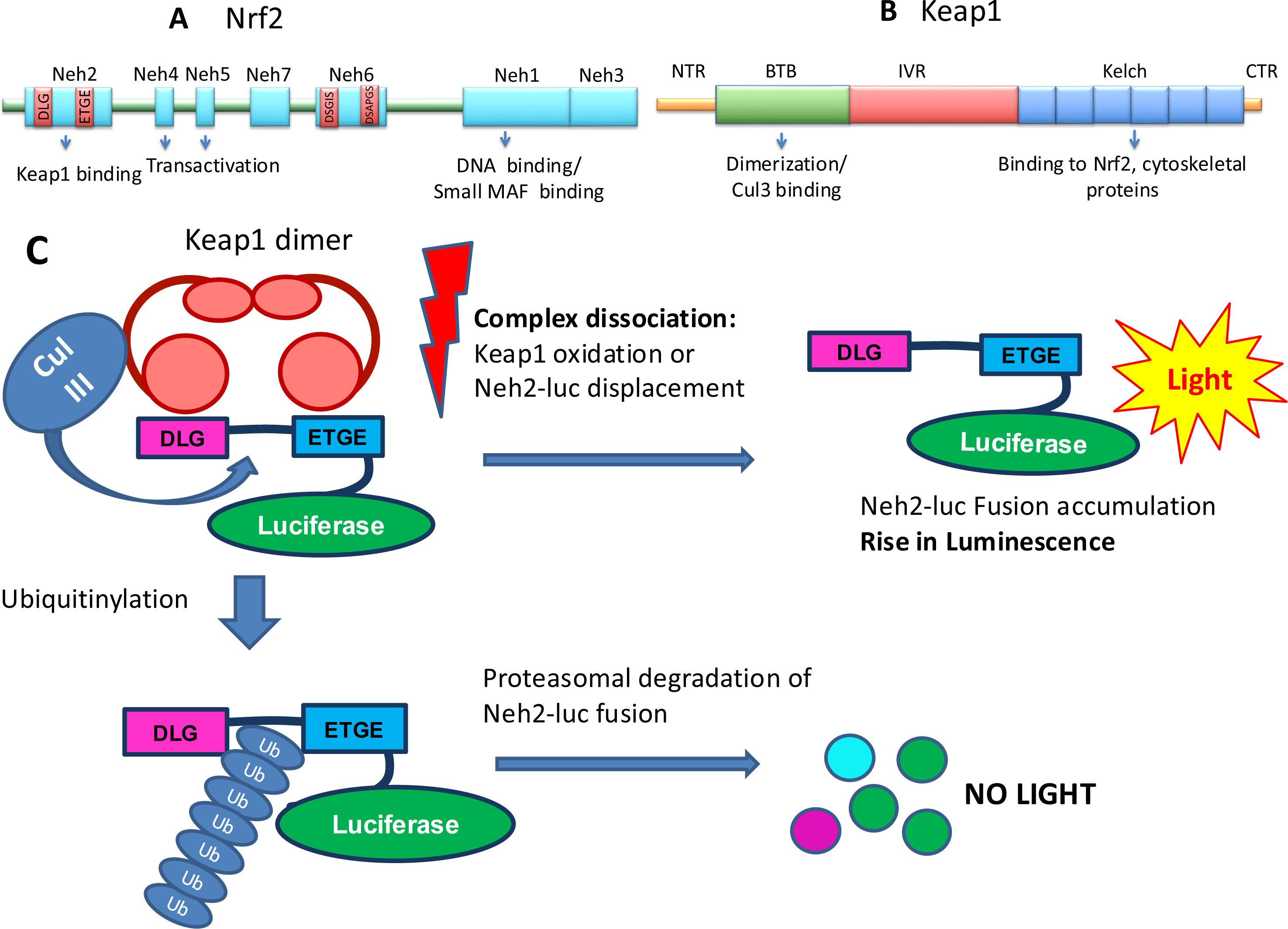
Domain structures of Nrf2 (A) and Keap1 (B), and mechanistic scheme of Neh2-luc reporter action (C). Neh2 in Nrf2 is the domain recognized by the Kelch domain in Keap1 through a specific interaction with the DLG and ETGE binding motifs in Neh2. Keap1 dimer is an adaptor protein in Cullin III ubiquitin ligase complex, which performs ubiquitinylation of Neh2 domain lysines to label Nrf2 or, in this case, Neh2-luciferase fusion protein for proteasomal degradation. Disassembling of Keap1-Cullin III-Nrf2 complex induced by covalent modification of Keap1 or by displacing Nrf2 results in stabilization of Nrf2 or, in the case of the reporter, Neh2-luc fusion protein, monitored by an increase in luminescence.

**Fig. 2.**
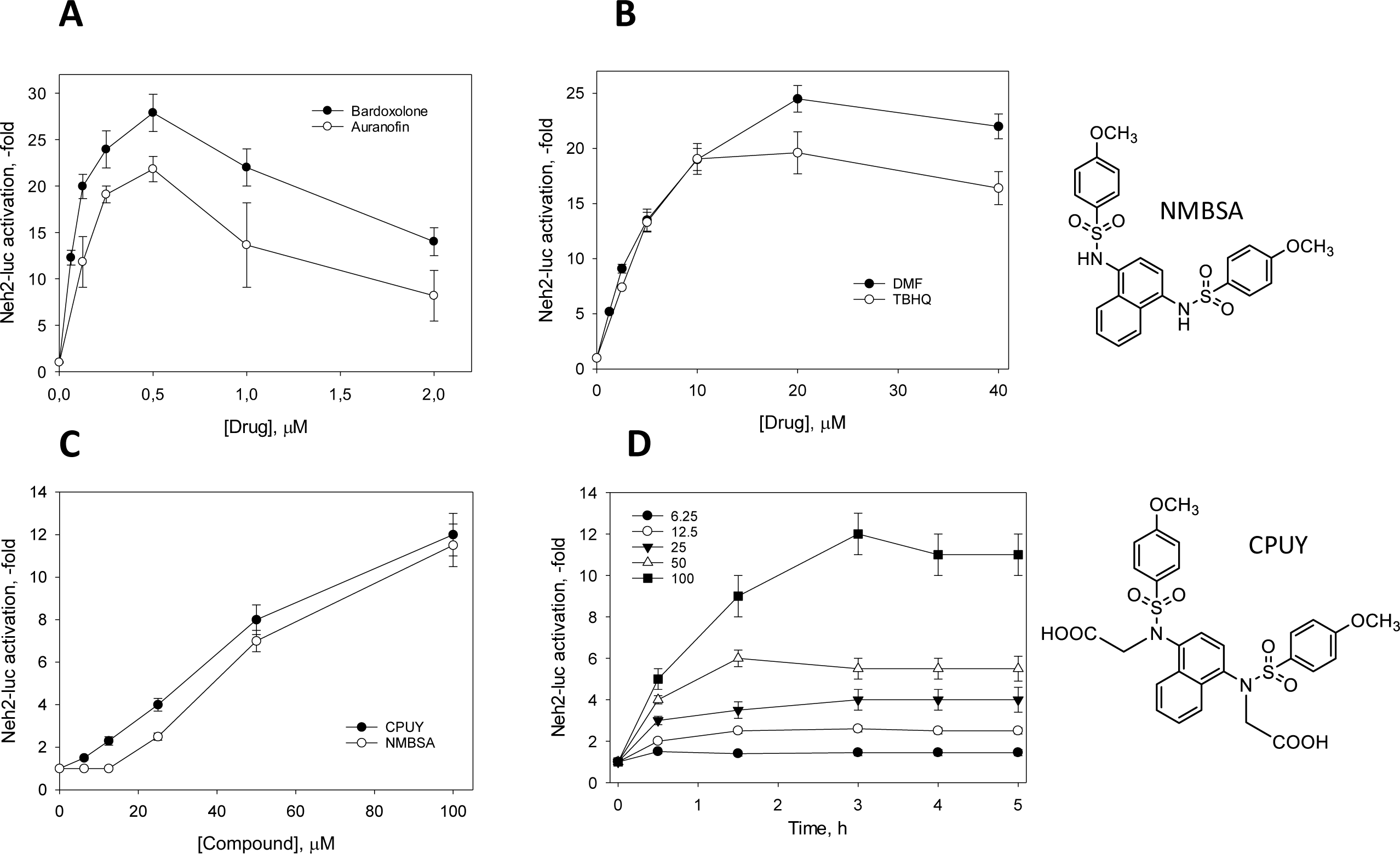
Activation of Neh2-luc reporter with various Nrf2 activators. Luminescence is normalized to the background luminescence in the absence of an inducer. A: bardoxolone and auranofin; B: Dimethyl fumarate (DMF) and *tert*-butylhydroquinone (TBHQ); C: Displacement activators CPUY and NMBSA; End-point assay at 3 h incubation. D: Time-course of reporter activation with NMBSA. Mean ± SEM.

True displacement activators such as CPUY exhibit a specific time course of Neh2-luc reporter activation with a characteristic plateau reached for each concentration of the displacement agent (Fig. 2D) at the time of system re-equilibration (Smirnova et al. 2011), which in this particular case is ca. 3 h. The kinetic behavior of Neh2-luc reporter in the presence and absence of displacement activators can be quantitatively described due to its similarity to the actual endogenous mechanism of Nrf2 degradation. The Neh2-luc fusion protein is generated with the M_0_ rate and then binds to free Keap1 protein (reaction 1 in Fig. 3), which is in equilibrium with Keap1-bound endogenous Nrf2 continuously produced with the X_o_ rate (reaction 2 in Fig. 3) and other client proteins (not shown in the scheme). In the presence of a displacement activator, the equilibrium between all Keap1-bound forms shifts towards the inhibitor-bound Keap1 (reaction 3 in Fig. 3). Hence, free Keap1 is less available for the binding of the luciferase fusion protein, leaving a higher steady-state concentration of the fusion protein. As calculated in (Khristichenko 2019), the reporter luminescence background corresponds to the steady-state concentration of all undegraded forms of luciferase fusion and is equal to 60 nM. The rate of luciferase fusion protein production – M_o_ – was determined from the maximum rate for alkylating activators as ca. 8 nM/min (Khristichenko 2019). At equilibrium, the rate of fusion protein production M_o_ must be equal to the rate of fusion protein degradation, which is linearly proportional to its steady-state concentration (60 nM). Hence, the first-order rate constant for the rate-limiting step of the reporter performance has been calculated as 8 nM/min divided by 60 nM, *k_lim_* = 2.2x10^-3^ s^-1^. This value is significantly lower than the rate constants reported for proteasomal degradation in the literature, which may reflect the binding and ubiquitination step in the Keap1-Cullin III complex. The Neh2-luc reporter validation studies (Smirnova et al. 2011) indicated that the rate-limiting step in Neh2-luc reporter degradation corresponds to Keap1 binding to the Neh2-luc fusion protein rather than proteasomal degradation. Therefore, the observed first-order rate constant *k_lim_* = 2.2x10^-3^ s^-1^ is close to the *k_1_*[Keap1] value, and if the value for the binding rate constant is known, free Keap1 concentration can be estimated.

**Fig. 3.**
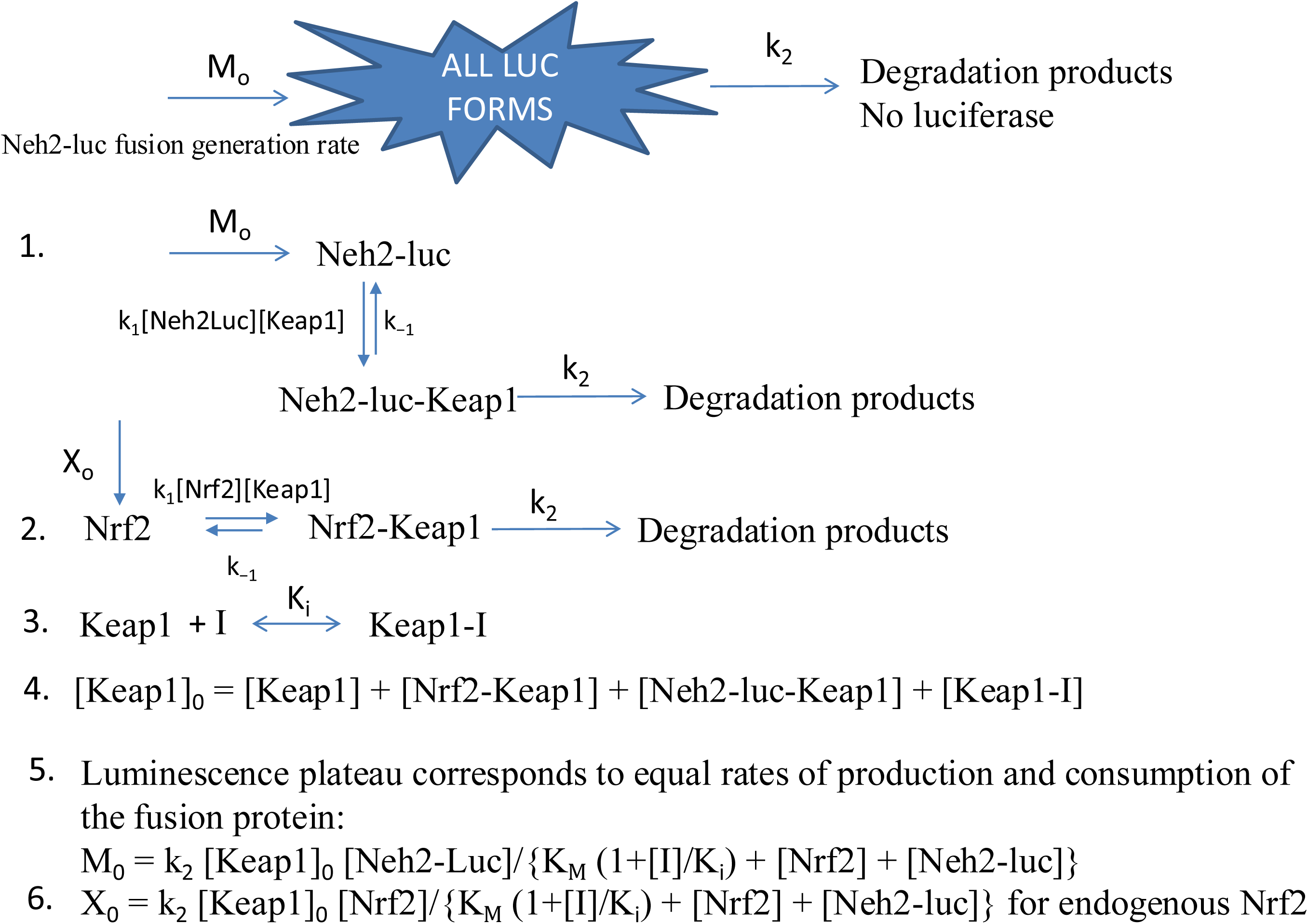
Kinetic scheme for Neh2-luc reporter action in the presence of a reversible inhibitor. Reaction 1: Neh2-luc fusion protein is continuously produced with *M_o_* rate; Neh2-luc fusion protein forms a complex with free Keap1 with the association rate constant *k_1_* and dissociation rate constant *k_-1_,* the following steps of ubiquitination and proteasomal degradation are shown as an overall step with *k_2_* rate constant. Reaction 2: Endogenous Nrf2 protein is synthesized continuously with *X_o_* rate and undergoes the same binding and transformation steps as the fusion Neh2-luc. Reaction 3: Reversible inhibitor I binds to free Keap1 with the inhibition constant *K_i_*. Equation 4: Material balance for all Keap1 forms. Equation 5: Equilibrated rates for production and consumption of Neh2-luc fusion protein in the presence of a reversible competitive inhibitor, *K_M_ = (k_2_ + k_-1_)/k_1_*. Equation 6: Equilibrated rates for production and consumption of endogenous Nrf2 protein in the presence of a reversible competitive inhibitor.

The Neh2 domain in the fusion protein gets stretched between the Kelch domains in the Keap1 dimer through the latter specific interaction with DLG and ETGE motifs in Neh2 to facilitate ubiquitination of Neh2 lysine residues by Cullin III ubiquitin ligase, leading to subsequent degradation of the Neh2-luc protein as a whole (Fig.1C). The second-order rate constants for equilibrium binding of DLG and ETGE motifs to Kelch domain were determined by surface plasmon resonance in (Fukutomi et al. 2014) as 6.1 x 10^4^ and 3.45x10^6^ M^-1^s^-1^, respectively. However, ETGE binds in two steps, and the second step is very slow – 1.3x10^-3^ M^-1^s^-1^, giving an association rate constant for ETGE equilibrium binding of about 4x10^3^ M^-1^s^-1^ (as a product of multiplication of two binding constants for two consecutive equilibrium steps). This estimate - *k_1_* = 4x10^3^ M^-1^s^-1^ - can be used as the maximum rate constant for Neh2-domain binding to Keap1 dimer, and knowing that the reporter’s rate-limiting step is mainly dependent on Neh2-luc fusion interaction with Keap1, one can calculate the intracellular Keap1 concentration by dividing the first order rate-limiting constant *k_lim_* = 2.2x10^-3^ s^-1^ to the second order rate constant of Nrf2 peptide interaction with Keap1 *k_1_* = 4x10^3^ M^-1^s^-1^ and get an estimate of minimum intracellular concentration of free Keap1 as 0.55 μM. This estimate points to one of possible reasons for the observed offset in the EC_50_ reported for displacement activators in the cell-based reporter assays – high, likely micromolar, concentrations of total intracellular Keap1, a versatile adaptor functioning as a trapping agent for multiple protein clients into the Cul III ubiquitin ligase complex. The amount of Keap1 protein clients competing in the cell with a displacement activator could have an additional effect on the biological potency of a displacement activator, depending on the absolute concentration of Keap1 client proteins and the activator’s K_D_ value. The reported K_D_ values for ETGE and DLG Nrf2 peptides are 28 and 130 nM (Fukutomi et al. 2014), respectively. Thus, to successfully compete with homogeneous Nrf2 binding, a displacement activator should have a K_D_ in the same range or lower. A third reason for the offset could be the non-specificity of Keap1 displacement activators, as there are at least 40 Kelch-containing proteins in the human genome, which we discussed in our recent review (Hushpulian et al. 2021).

### 3.2. High intracellular concentrations of Keap1 limit the effectiveness of displacement activators

The absolute amounts of intracellular Keap1 and Nrf2 will have a profound effect on the biological potency of Nrf2 displacement activators. Therefore, Nrf2 and Keap1 quantification have been performed for two cell lines, e.g. the untransformed neuroblastoma SH-SY5Y line and its transformed variant that stably expresses Neh2-luc fusion. Since the Neh2-luc fusion protein competes with an endogenous Nrf2 protein, Nrf2 concentration might be higher in the transformed cell line than in the untransformed line, which could lead to partial activation of the Nrf2 genetic program in the reporter line, and therefore, a higher level of Keap1 protein due to its feedback regulation – Keap1 gene is a known Nrf2-target.

A representative blot to determine the absolute amounts of Keap1 and Nrf2 proteins in the cell lines using recombinant Keap1 and Nrf2 proteins, shown in Fig. 4A, clearly demonstrates that expression of the Neh2-luc fusion protein in the cell line results in an increase in both Nrf2 and Keap1 protein levels. The endogenous Nrf2 is spared from degradation due to the constitutive synthesis of the fusion, which competes for Keap1 binding. Consequently, Nrf2 upregulates Keap1 through a feedback mechanism. The levels of Keap1 and Nrf2 proteins are ca. 5-fold and 2.5-fold higher, respectively, in the Neh2-luc reporter cell line compared to the non-transformed cell line (Fig.4 A, B). The Nrf2 content per 1,000 cell is close to 0.1 fmol in the untransformed line and 0.25 fmol in the reporter line (from the blot in Fig. 4A). Recalculated into absolute amounts, these numbers are equal to 60,000 and 150,000 molecules, respectively, close to those previously reported for several cancer cell lines, where the absolute values for Nrf2 varied from 49,000 to 190,000 molecules per cell (Iso et al. 2016). Taking 12 μm diameter and 3 μm thickness for the neuroblastoma cells in the monolayer, the cell’s estimated volume is ca. 340 μm^3^. Using this number, the intracellular concentrations of Nrf2 are calculated as ca. 0.30 μM and 0.75 μM for the plain and transformed cell lines, respectively. For Keap1 protein, the content in the plain cell line of ca. 0.9 fmol per 1,000 cell (540,000 molecules per cell) is increased up to 3.8 fmol (almost 2 million molecules per cell), or from ca. 2.5 μM to 12 μM (Fig. 4A, B). The number of Keap1 molecules in the non-transformed neuroblastoma cell exceeds those reported for several cancer cell lines, where the absolute values for Keap1 varied from 45,000 to 300,000 molecules. However, the estimated Keap1 cytosolic concentration in RAW264.7 cells, calculated as 1 μM, (Iso et al. 2016) is not far from the 2.5 μM determined in this study. In the same study (Iso et al. 2016), the absolute values for the Nrf2 and Keap1 protein content have been measured after cell treatment with diethyl maleate, an electrophilic Nrf2 activator, a Michael acceptor that covalently modifies Keap1 cysteines: within three hours of treatment no change in Keap1 amount was observed whereas Nrf2 protein could rise and reach concentrations from 330,000 to 710,000 molecules per cell (Iso et al. 2016).

**Fig. 4.**
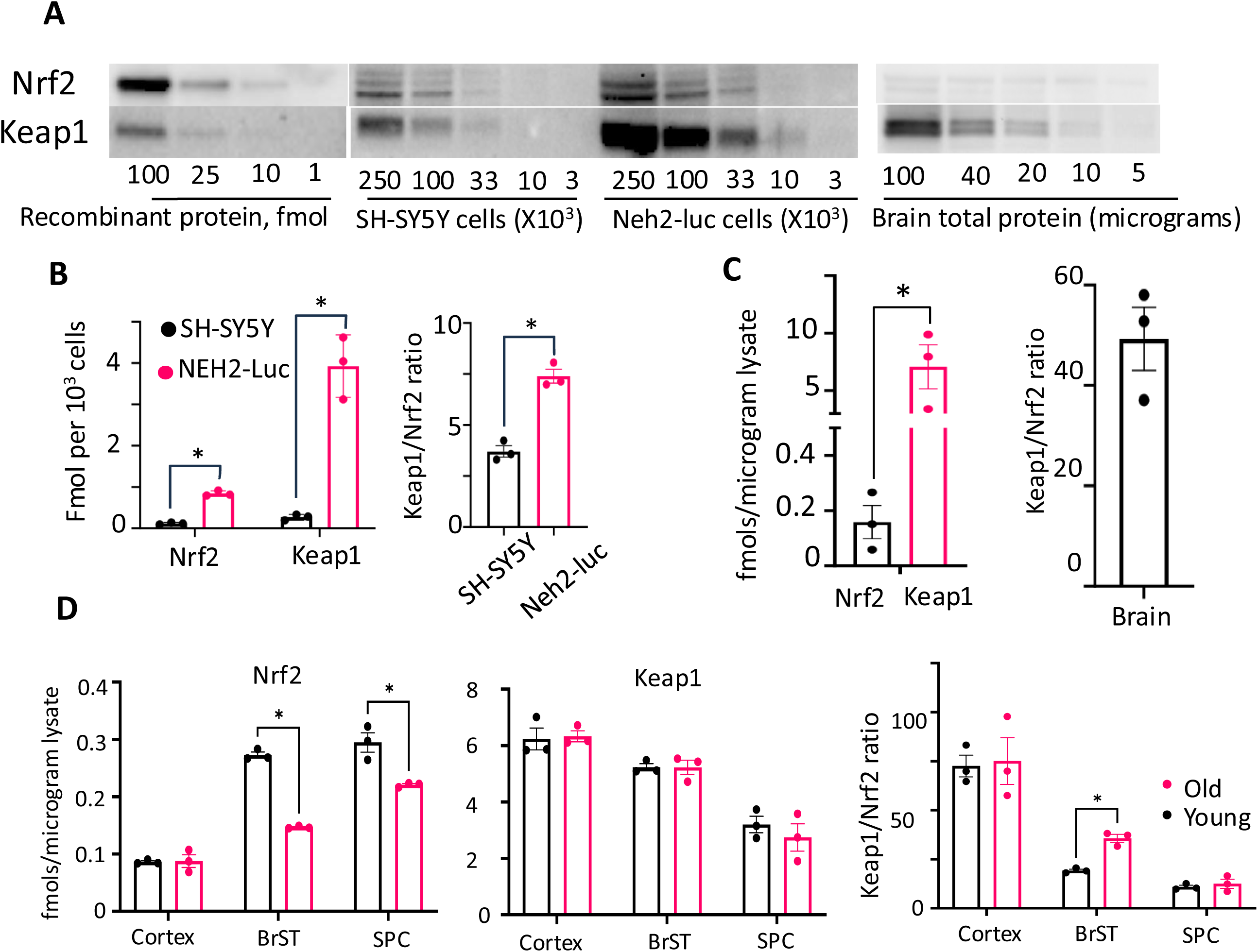
Nrf2 and Keap1 protein content and their ratios in neuroblastoma (SH-SY5Y), Neh2-luc reporter lines, in the mouse whole brain, cortex, brainstem, and spinal cord. Quantification of Nrf2 and Keap1 protein content and ratio in the untransformed neuroblastoma cell line (SH-SY5Y) and Neh2-luc reporter line (A, B), the mouse whole brain (A, C), and in the cortex, brainstem (BrST), and spinal cord (SPC) of young and old animals (D). Mean ± SEM.

The Keap1/Nrf2 ratio in the Neh2-luc reporter cells doubles compared to the non-transformed neuroblastoma cell lines and reaches ca. 7-8 fold (Fig. 4B). Even if one recalculates these numbers into the Keap1 dimer concentration, the latter is still much higher than that of endogenous Nrf2. The significant excess of Keap1 explains the effective degradation of the constitutively produced Neh2-luc fusion, resulting in a steady-state concentration of 60 nM. Given the K_D_ values for Nrf2 binding motifs to Kelch within the 30-130 nM range (Fukutomi et al. 2014), both Nrf2 and Neh2-luc fusion proteins mainly exist in a complex with Keap1. Concerning Nrf2 displacement activators, besides the high Keap1 content, the challenge lies in the need to displace both endogenous Nrf2 and Neh2-fusion from their complexes with Keap1. Irreversible alkylating activators will react with available Keap1 thiols stoichiometrically, so we may see the reporter activation with their nanomolar concentrations like in Fig. 2A. In contrast, true displacement activators could be active only in high micromolar concentrations because they need to bind most of Keap1 protein reversibly and thus, outcompete the endogenous Nrf2, whose concentration is continuously growing upon Keap1 re-equilibration with the displacement activator. One may even estimate the rate of Nrf2 production in the cell (X_0_ in reaction 2 in Fig. 3), based on the amount of Nrf2 build-up amount in various cell lines upon Keap1 irreversible modification reported in (Iso et al. 2016): 350,000 to 500,000 molecules per cell in 3 h, which corresponds to 15.6 -22.2 nM/min rate for Nrf2 production. The maximum rate for Neh2-luc expression activated by CPUY estimated from Fig. 1D as ca. 8-10 nM/min is 2-3-fold less than the expression rate of endogenous Nrf2 production calculated from published data (Iso et al. 2016).

Assuming the same transformation scheme and equal rate constants for transformations of endogenous Nrf2 and Neh2-luc fusion, the ratio of their production rates should be close to the ratio of their steady-state concentrations (see Eq.5 and 6 in Fig. 3). In the presence of an inhibitor, if the luminescence plateau at EC_50_ (for example, at 50 μM CPUY in Fig. 2D) corresponds to a 5-fold increase over the background level of 60 nM, the same increase in the concentration of endogenous Nrf2 may be expected. Thus, the total concentration of free ([Nrf2] + [Neh2luc]) will be around 4 μM. This means that to successfully compete with high intracellular concentrations of Nrf2 and Neh2-luc fusion, the value for *K_M_ (1+[I]/K_i_)* must exceed their overall concentration. If we estimate the *K_M_* value close to the reported K_D_ values (28 nM and 140 nM), the inhibitor should be at least 20-100 times greater than its *K_i_* value. Again, this is a rough estimate to explain the consequences of competition with endogenous Nrf2 for the biological potency of a displacement activator in the Neh2-luc fusion assay. In addition to Nrf2, Keap1 has multiple client proteins, whose concentration may also rise and exert additional competition against the displacement activator (see the list of 14 Keap1 clients in our review (Hushpulian et al. 2021). Therefore, Nrf2 displacement activators will never reach the maximum activation threshold for irreversible Nrf2 activators in the Neh2-luc assay, as shown in Fig. 2A and B (ca. 25-fold activation). In contrast, NMBSA and CPUY exhibit a 12-fold activation level, corresponding to the re-equilibrated concentration of the fusion protein (Fig. 2C and D).

Given the absolute amounts of Keap1 and Nrf2 in cancerous cell lines and high rates of Nrf2 production upon Keap1 inhibition, high-throughput screening for Nrf2 displacement activators in cell-based assays should be performed at high micromolar concentrations of drugs (20-100 μM). In other words, if a prospective displacement activator shows EC_50_ below 1 μM in the cell-based reporter assay, an irreversible modification of Keap1 can be suspected. True, reversible displacement activators will exhibit high micromolar EC_50_ values in cell-based assays due to the aforementioned issues, both for fusion reporters and transcription reporters, such as ARE-luc. For example, a displacement activator identified and described in (Hu et al. 2013) shows K_D_ in homogeneous assay close to 1 μM and EC_50_ in ARE-luc assays of ca 12-18 μM, NMBSA exhibits cellular activity within 10-100 μM range (Lu et al. 2016) and CPUY within 1-100 μM (Lu et al. 2016). An asymmetric NMBSA-type compound, RA839 (see Fig.S1), showing K_D_ of 14 nM in FP assay, exhibited EC_50_ of ca. 40 μM in the cell-based ARE-luc assay (Winkel et al. 2015). A CPUY analog – compound K22 - with a tetramethylbenzene scaffold instead of a naphthalene (Sun et al. 2022) with K_D_=180 nM induced the expression of Nrf2 genes at concentrations of 5-10 μM. In our earlier work on aspirin-containing prodrugs, an Nrf2 displacement activator with a naphthalene core, like NMBSA, and asymmetric aspirin-containing “arms” was shown to behave as a potent displacement Nrf2 activator (EC_50_ of ca 10 μM in the reporter assay) due to partial hydrolysis catalyzed by the cell esterase, leading to a metabolite perfectly fitted into the Kelch domain - with C-docker energy twice as good as that for NMBSA (Gaisina et al. 2021).

To evaluate the prospects of genuine true displacement activators for *in vivo* use, one must determine the absolute amounts of Keap1 and its major client protein, Nrf2, in tissues where oxidative stress is considered a key damaging factor and where the activation of the Nrf2 genetic program could be a “magic bullet.” One of the most sought-after applications of Nrf2 activators is their potential to mitigate oxidative stress in the brain, thereby promoting healthy aging and counteracting age-related cognitive decline. As shown in Fig. 4A & C, the content of Nrf2 in brain is very low as compared to that of Keap1, almost 50 times lower (Fig. 4C). The amount of Nrf2 varies from 0.1 to 0.2 fmol per μg protein (Fig. 4C left panel), which is ca. 4-5 times lower than the value reported for the rat brain in (Edres et al. 2021) as 0.8 μg/mg protein. To recalculate the values presented in Fig. 4C-right panel & D into the concentration units, one may take the protein content estimate as 0.2 g/mL (Milo 2013), and, thus, end up with ca. 0.02 μM Nrf2 and 1 μM Keap1 in the brain. The absolute amounts of Keap1 and Nrf2 proteins determined in young and old mice for three regions – cortex, brainstem, and spinal cord – are shown in Fig. 4D. No significant changes in Nrf2 and Keap1 content are observed in the cortex with aging (Fig. 4D). Interestingly, Nrf2 protein content were significantly reduced in the brainstem and spinal cord in the aged mice compared to younger mice (Fig 4D). However, Keap1 protein levels showed no difference between young and old mice in the brainstem and spinal cord (Fig. 4D). Thus, these changes in Nrf2 and Keap1 protein content as a function of age resulted in a significant increase in Keap1/Nrf2 ratio in the brainstem from older mice (Fig. 4D). Notably, early involvement of brainstem structures in the disease process has been described for PD and AD (Grinberg, Rueb, and Heinsen 2011). Neuropathological studies reveal that the tau pathology in prodromal and preclinical AD may originate in the brainstem nuclei (Dutt et al. 2021); the brainstem is a region that is compromised in early PD (Vijayan et al. 2020).

In all cases, a significant excess of Keap1 over Nrf2 in the brain (Fig. 4) opens an opportunity to upregulate the Nrf2 genetic program by targeting Keap1. However, for a displacement activator, there will be a need to deliver high micromolar concentrations into the brain. In contrast, irreversible Nrf2 activators that are specific only for Keap1 will be required in amounts matching those for Keap1-active thiols (ca. 10 nmol Keap1 in a 2 g brain). Under the most optimistic scenario, not accounting for the presence of multiple protein clients of Keap1 in the cell, one may expect an order of magnitude higher amounts of an Nrf2 displacement activator to be comparable to a hypothetical Keap1-specific irreversible inhibitor. In real life, the dose of an Nrf2 displacement activator is expected to be significantly higher, as the accumulation of endogenous Keap1 clients will competitively interfere with the drug’s action.

In the recent work, CPUY molecule was modified by replacement of both carboxy-groups with amides and one of the *p-*methoxyphenyl substituents with a *p-*aminophenyl group to generate 2-((*N*-(4-((4-amino-*N*-(2-amino-2-oxoethyl)phenyl)sulfonamido)naphthalen-1-yl)-4-methoxyphenyl)sulfonamido)acetamide (coded as NXPZ-2), a displacement activator with K_D_ of 94 nM and an improved ability to cross the BBB (Sun et al. 2020). The compound ameliorated learning and memory dysfunction in Aβ1-42-treated mice at doses ranging from 52 to 210 mg/kg (Sun et al. 2020). These doses are higher than those typically used for dimethyl fumarate (DMF), an FDA-approved Nrf2 activator of an alkylating nature, in various scenarios of neurodegenerative diseases (Majkutewicz 2022). However, NXPZ-2 showed benefits in the post-treatment scenario, with the drug administration starting 7 days after the Aβ1-42 injection into the hippocampus (Sun et al. 2020). In contrast, DMF and other alkylating Nrf2 activators are usually effective in a pretreatment regimen. Modifying the amine group in NXPZ-2 with a phosphodiester substitution (novel variant named POZL) permitted the effective dose to be lower than 40 mg/kg (Sun et al. 2023). However, the phosphodiester substitution is an active group, so the variant generated cannot be considered a pure displacement one, especially given that the binding affinity for POZL for Keap1 is four times lower than that of NXPZ-2 (Sun et al. 2023). So, POZL likely works through “suicide” displacement (i.e., a displacement followed by covalent modification).

To develop a potent *displacement* activator, structural optimization aimed at lowering the K_D_ to single-nanomolar values is insufficient because the intracellular Keap1 content is high. Another concern is that displacement activators could be non-specific by targeting all Kelch-type proteins. From this perspective, oxidative alkylating agents are more specific for Keap1, as it serves as a redox sensor and thus differs from many other Kelch domain proteins. This dictates a different strategy for designing effective Nrf2 displacement activators intended for *in vivo* application – a combination of an Nrf2 displacement scaffold or peptide with an alkylating or pro-oxidative motif in a single molecule.

### 3.3. Targeting Keap1 with Nrf2 displacement activators bearing an alkylating motif

NMBSA has a *symmetric* structure with two sulfonamide “arms.” If one of the arms is removed and the naphthalene moiety is replaced with a quinoline, molecules like 8TQ and 8QBSA (Fig. 6) may also function as displacement activators. The quinoline scaffold preserves the affinity for the Kelch domain, and 8QBSA demonstrates a reasonable value for C-docker energy (– 33 kJ/mol) in accordance with modeling studies. 8-Aminquinoline, by itself, exhibits some chemical activity compared to the inert naphthalene scaffold. The introduction of chloro- or bromo-substitution instead of the second “arm” in the 5^th^ position creates a strong alkylating molecule preserving the ability to target Keap1 (Fig. 6B). Comparison of the 5-chloro-substituted analog 5-Cl-8QBSA with the 5-bromo-substituted analog 5-Br-8QBSA revealed that the presence of the larger halogen slightly reduces C-docker energy from -39 to -38 kJ/mol, respectively, but the energies are much closer to C-docker energy for NMBSA (-48 kJ/mol), used for validation of the docking procedure (the docking model should overlap with the actual position of NMBSA in the crystal structure). Of note, the absence of the second sulfonamide “arm” permits a deeper positioning of 5-halogenated analogs of 8-quinolinyl-benzenesulfonamide (8QBSA) (shown in purple) inside the Kelch pocket compared to NMBSA (shown in red in Fig. 6A). However, the introduction of a potent alkylating motif leads to unavoidable toxic side effects due to non-specific alkylation of cellular components on the way to the target, and the activation profile shows a peak characteristic for the onset of toxicity (Fig. 6B). Many alkylating activators of Nrf2 like natural ones (sulforaphane, myricetin, mangiferin, isoastilbin, quercetin) as well as synthetic ones like dimethyl fumarate and bardoxolone are reported to play a protective role against different diseases associated with inflammation. However, the exact mechanistic link between Nrf2 activation and its anti-inflammatory effects requires further research (Pant et al. 2024). 4-Methyl-*N*-(quinoline-8-yl)benzenesulfonamide, also known as 8-(tosylamino)quinoline (8-TQ), a mild Nrf2 activator similar to its 2,4,6-trimethyl-analog 8QBSA (Fig. 6B), has been demonstrated as an effective anti-inflammatory drug, which alleviated the signs of LPS-induced hepatitis in mice when administered at 20-40 mg/kg doses for 3 days (Jung et al. 2012). However, the 8-aminoquinoline scaffold-derived drugs, despite their well-known use as antiparasitic drugs, are also known for serious side effects and are not a perfect choice for “benign scaffold” Nrf2 activators.

**Fig. 5.**
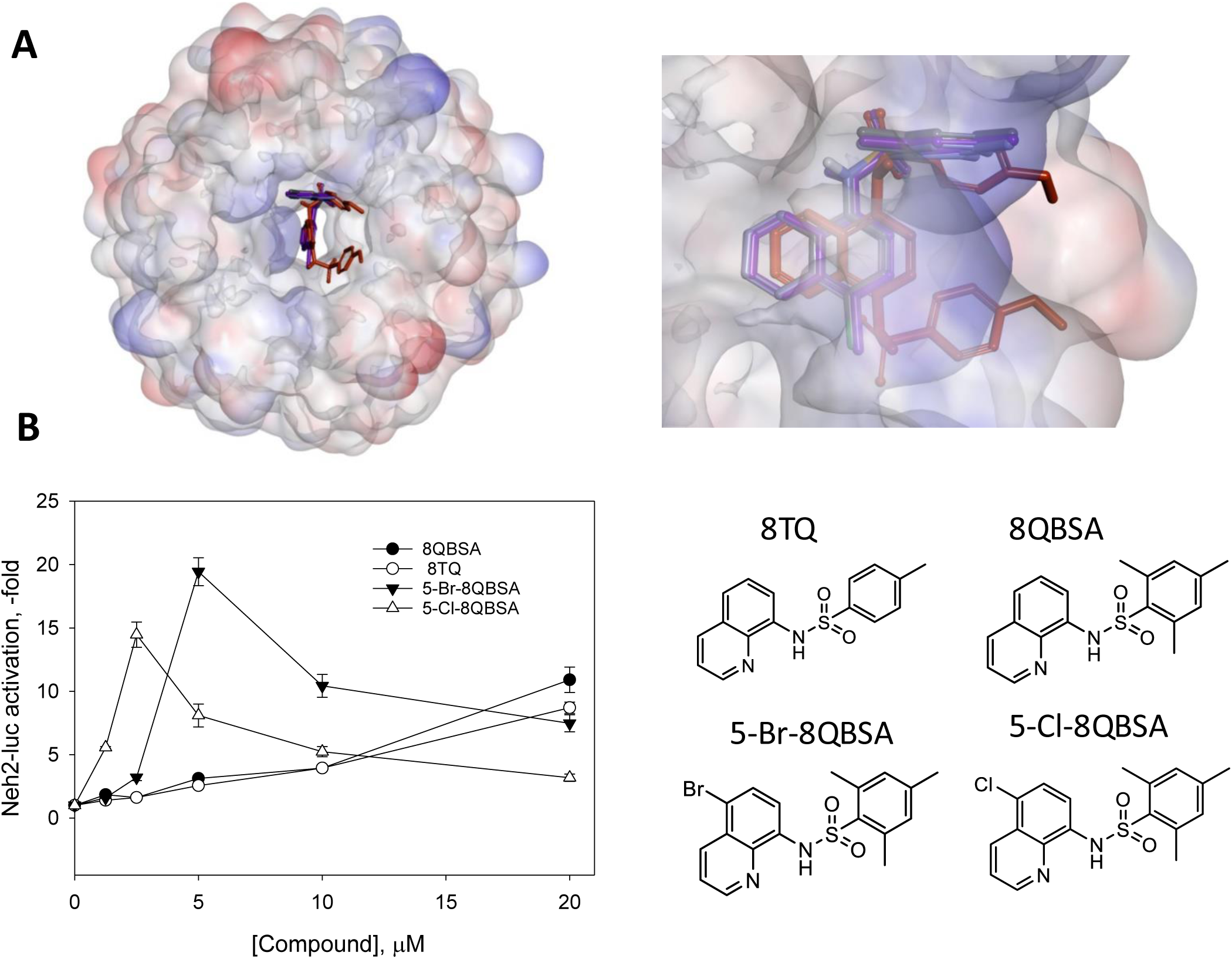
Aminoquinoline activators of Nrf2. A: docking into Keap1 Kelch domain. B: Neh2-luc reporter activation at 3 h incubation. The structures of pure displacement 8TQ and 8QBSA, and alkylating displacement variants with halogen substitution in the 5^th^ position – 5-Cl-8QBSA and 5-Br-8QBSA – are shown.

**Fig. 6.**
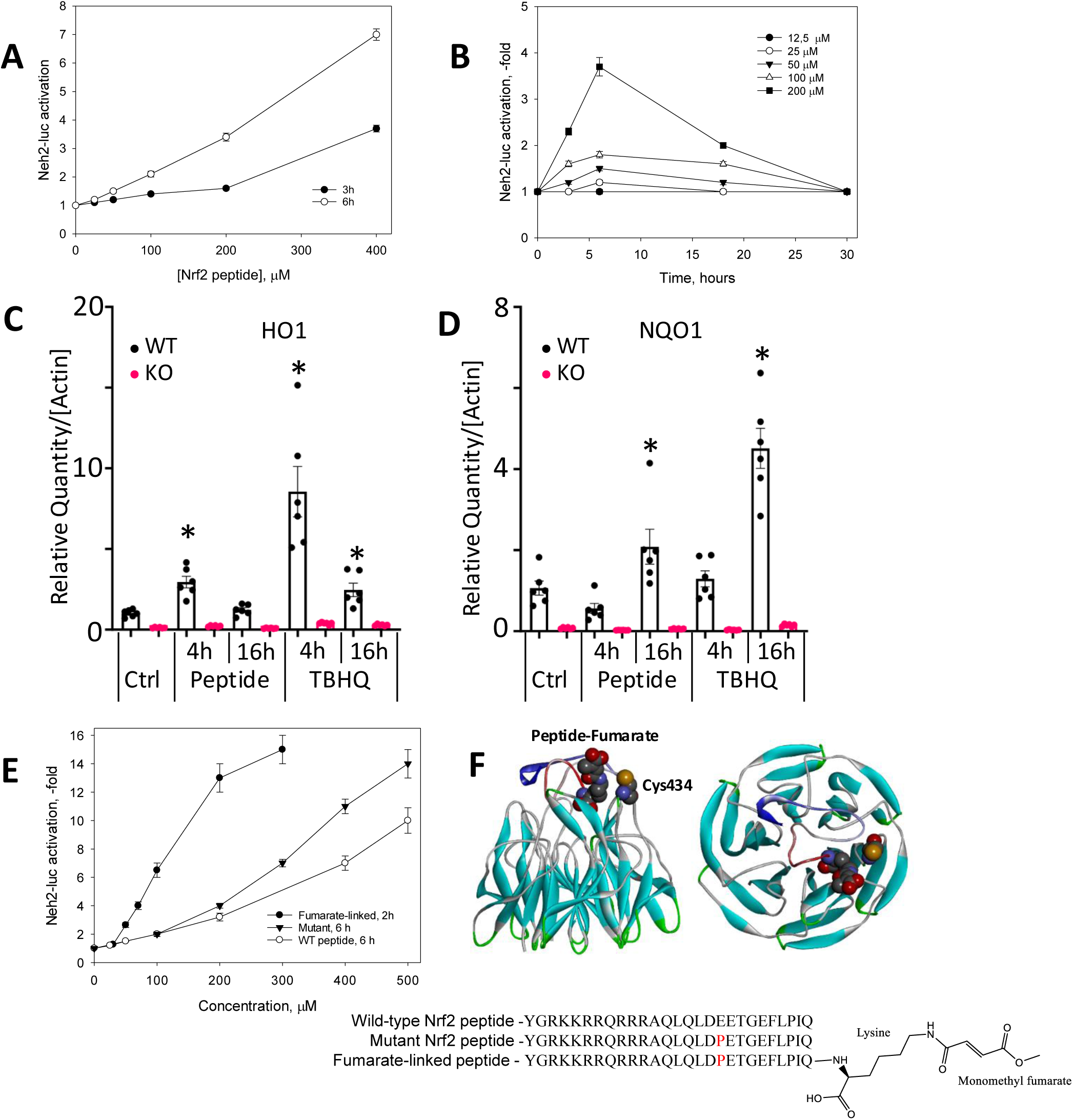
Nrf2 activation with cell-permeable Nrf2 peptides. End-point (A) and time course (B) titration for the wild-type and Glu->Pro mutant peptide; RT-PCR for Nrf2 target genes heme oxygenase 1 HO1 (C) and NQO1 (D) induced by 200 μM wild-type Nrf2 peptide at 4 and 16 h incubation in wild-type and Nrf2 KO mouse embryonic fibroblast in comparison to the action of 20 μM TBHQ; comparison of fumarate linked mutant Nrf2 peptide to the wild-type and mutant peptide without any alkylation motifs in Neh2-luc assay (E), and docking of fumarate-linked peptide into the Kelch domain showing Cys434 in close proximity to the fumarate tail (F).

Instead, by making asymmetric NMBSA type compounds with a 4-amino-1-naphthol based scaffold, the chemical reactivity can be increased due to the second “asymmetric” arm with a pro-oxidative motif, as exemplified by derivatives S47 (Zhuang et al. 2014) and 20c (Lu et al. 2020) (structures shown in Fig. S1). Both molecules exhibit the properties of alkylating displacement activators providing targeted chemical modification of Keap1: S47 with K_D_ of 1.9 μM is active at 4 μM in the cell (Zhuang et al. 2014). The recently characterized compound ZJ01 features a coumarin scaffold instead of a naphthalene one and additionally bears substitutions with obvious alkylating potential (Fig. S1). Although ZJ01 affinity for Keap1 drops (K_D_ is only 5.1 μM in FP assay), this compound exhibits an EC_50_ value of 8 μM *in vitro* in the cell-based assays, and is active *in vivo* at the 5-10 mg/kg doses in LPS-inflammation model on mice (Jiang et al. 2018), thus providing strong evidence for the engagement of alkylating mechanisms.

### 3.4. Cell-permeable Nrf2 peptides and their optimization

Small-molecule displacement activators are chemical compounds that could exert undesirable side effects due to the chemical or enzymatic transformation of their structural core and substituents within the cell. To model a benign, “natural” approach to target Keap1, we used cell-permeable Nrf2 Peptides by linking the “ETGE” binding motif to an N-terminal TAT peptide. In this work, a cell-permeable peptide construct involving a 16-mer “ETGE” binding motif attached to the TAT-sequence, named as a wild-type Nrf2 peptide with the sequence YGRKKRRQRRRAQLQLDE**ETGE**FLPIQ, was active in the Neh2-luc assay only above 50 μM: the reporter activation slowly develops and peaks at 6 h incubation and then declines, likely due to peptide degradation (Fig. 7A, B). The longer time necessary to equilibrate the reporter compared to small-molecule displacement activators may reflect the two-step binding of the ETGE sequence that we mentioned earlier. The EC_50_ value in the reporter assay (ca. 200 μM) is orders of magnitude higher than the K_D_ estimate from the FP assay, and activation of gene expression by the peptide at 200 μM is much less than that for a covalent activator, TBHQ, used at 20 μM (Fig. 7C, D). The cell-permeable Nrf2 peptide triggers the Nrf2 genetic program only in cells expressing Nrf2, but not in Nrf2 KO mouse embryonic fibroblasts (MEF) (Fig. 7C, D).

**Fig. 7.**
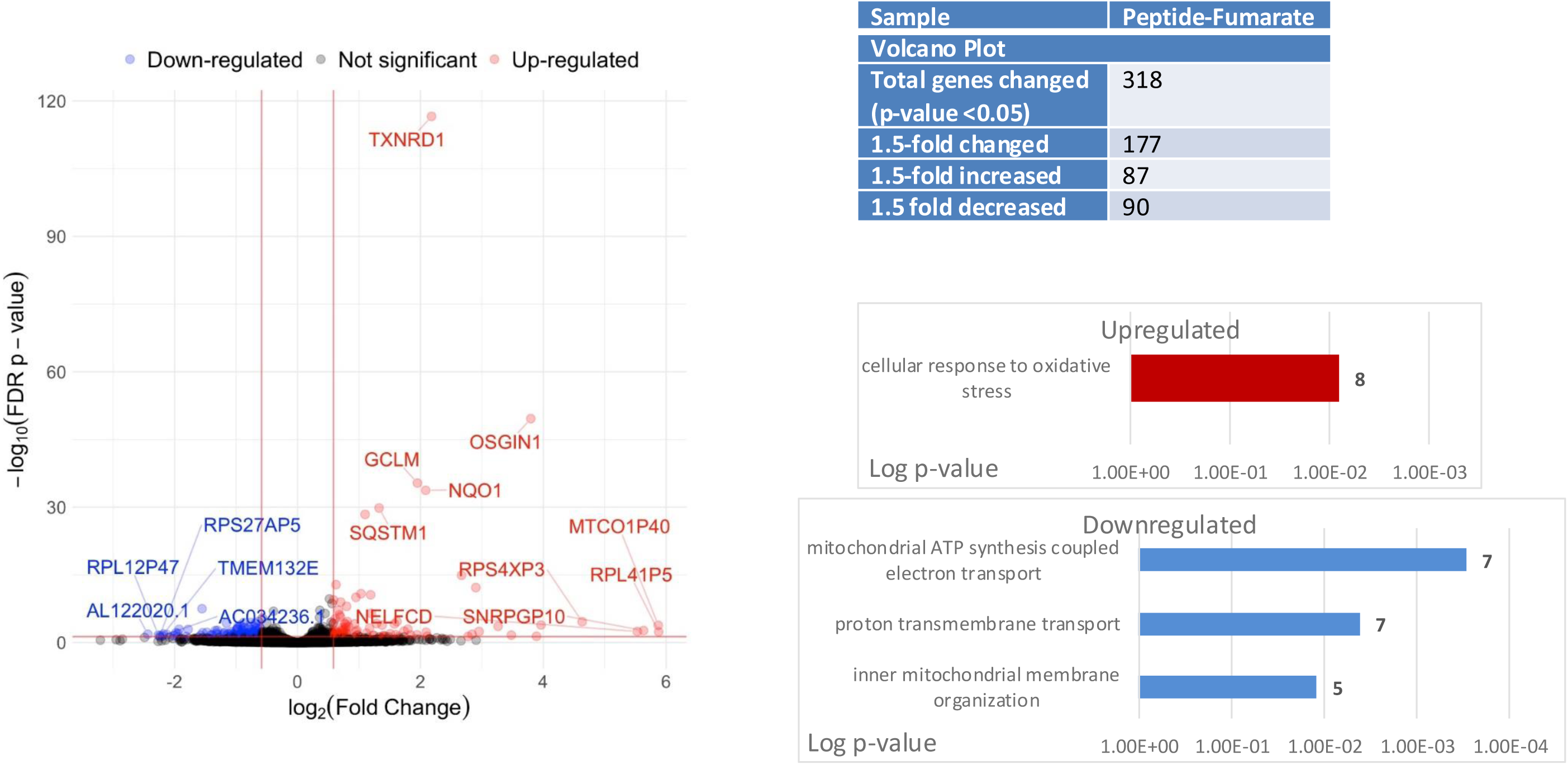
Volcano plot and affected GO Biological Processes in neuroblastoma cells treated with 100 μM cell-permeable fumarate-linked Nrf2 peptide at 5 h incubation.

Notably, HO-1 is recognized as a rapidly responding gene, whereas NQO-1 is a slowly responding target gene of Nrf2. Such an offset in EC_50_ for an Nrf2 peptide may indicate significant competition from the build-up of endogenous Nrf2, in addition to the Neh2-luc fusion protein itself, which likely exhibits a much higher affinity for Keap1 than the TAT-Nrf2 Peptide. One may expect that mutations in the ETGE motif may improve the potency of the Nrf2 peptides based on the mutational analysis tested in the FP-assay: introducing proline ahead of the ETGE sequence instead of glutamic acid –DEETGEQ mutated to DPETGEL or DPETGEI, to match the sequence in p62, was reported to improve the K_D_ value by 50-fold (Hancock et al. 2012). However, mutation of glutamic acid to proline right ahead of the ETGE sequence in the cell-permeable Nrf2 peptide – YGRKKRRQRRRAQLQLD**PETGE**FLPIQ - showed a modest 2-3-fold improvement in EC_50_ in Neh2-luc assay (Fig. 7E). The values reported in the literature for the cell-permeable 14-mer TAT-Nrf2 peptide are K_D_ = 22 nM and EC_50_ > 40 μM for HO-1 induction (Steel et al. 2012), demonstrating a similar three orders of magnitude offset. This offset highlights the issue of reversible competition with the endogenous Nrf2, which is likely to exhibit a significantly higher affinity for Keap1 in the cell than in the FP assay.

We must note that there are two Kelch sites in the Keap1 dimer, and endogenous Nrf2 binds to both Kelch sites cooperatively. Stated another way, the local concentration of Nrf2 after dissociation from one of the two Kelch sites in the Keap1 dimer is high compared to the free concentration of a “displacement” activator. Because of this, the mean complex lifetime for Nrf2 will be much longer than the mean complex lifetime of univalent “displacement” activators. A recently reported multivalent “displacement” activator employs a synthetic, lipophilic polymer backbone with the LDPETGEFLRRRR peptide on each monomer unit (Carrow et al. 2024). An increase in local concentration of the Nrf2 peptide comes at the expense of its high molecular weight (10 to 15 kDa, depending on the degree of polymerization). Again, these polymers, which contain repeating Nrf2 peptide fragments, exhibit a sub-nanomolar affinity for the Kelch domain in an FP assay, but yield micromolar values in the ARE-luc assay in HepG2 cells. The values of EC_50_, being recalculated for the concentration of the peptide, give EC_50_ of 60 to 100 μM per peptide link for low (11 monomers) and high (27 monomers) polymerization degree variants, respectively (see Fig. S28B in (Carrow et al. 2024), just a slightly better values than those observed in Fig. 7A & B.

Two other issues limit the use of peptides, including the problem of low permeability through the cell membrane and instability within the cell. In the above paper (Carrow et al. 2024), with the help of a properly modified peptide sequence (quadrupole arginine residues at the C-terminus of the Nrf2 peptide) and polymer backbone, these issues were resolved, and the subsequent use of the polymeric Nrf2 peptide as a single intravenous injection demonstrated a promise in the post-treatment of myocardial infarction in a rat model (Mesfin et al. 2025). Despite the problems associated with the use of peptides, *in vivo* studies with Nrf2-peptides show that such treatment is truly benign and effective in post-treatment regimes. DEETGE-CAL-Tat peptide (NH2-RKKRRQRRR-PLFAER-LDEETGEFLP-CONH2) (Tu et al. 2015) in a rat model of global cerebral ischemia demonstrated a robust neuroprotection and preservation of hippocampal-dependent cognitive function, both at doses 30-100 μg in 5 μl of 0.9% saline injected by unilateral intracerebroventricular administration 30 min before ischemia and at doses 1-2 mg in 100 μl of saline (120 and 240 μg/d) 1–9 days post-treatment, beginning at 1 d after reperfusion.

Given the reversible mechanism of Nrf2 peptide action, linking the peptide to a chemical moiety with an alkylating or pro-oxidative motif may create an irreversible Keap1-targeted Nrf2 activator. To minimize side effects, the alkylation agent has yet to be activated or released from the peptide in the cell. A good candidate for such a mechanism could be monomethyl fumarate: a methyl fumarate-bound cell-permeable Nrf2 peptide synthesized by the addition of a lysine residue to the C-terminal of the mutant Nrf2 peptide via an amide bond (see structure in Fig. 6F). The nature of substitutions at the fumarate carboxy group strongly affects its cell-permeability and alkylating potency, with monomethyl fumarate (MMF) exhibiting the lowest potency in the cell-based assays (MMF<<DMF<diethyl fumarate<bis-salicyl fumarate) (Ahuja et al. 2016). Peptide-bound methyl fumarate will be slowly hydrolyzed to become an active alkylating agent in the cell. In this way, it may have sufficient time to target Keap1 before turning into a potent alkylating agent. Hence, the off-target effects of fumarate will be minimized. As shown in Fig. 6E, the fumarate-linked Nrf2 peptide exhibits a lower EC_50_; more importantly, it reaches the plateau of maximum activation within two hours. Computer modeling points to Cys 434 residue neighbors to C-terminal fumarate in the designed Nrf2 peptide (Fig. 6F). Therefore, there is a potential alkylation target in the Kelch domain right at the Nrf2 peptide binding site. If a fumarate-linked Nrf2 peptide functions as a Keap1-specific alkylating agent, one would expect it to activate the Nrf2 genetic program selectively, i.e., it would behave as a perfect Nrf2 activator.

### 3.5. Transcriptomic analysis of fumarate-linked Nrf2-peptide identifies “pure” Nrf2 target genes

We performed a transcriptomic analysis to investigate the specificity of the Nrf2-peptide in SH-SY5Y cells. We found that GO analysis identifies only one biological process activated by the Nrf2-fumarate peptide (Fig. 7) – “cellular response to oxidative stress.” The list of top-activated genes is presented in Table 1, and the genes can be divided into four subgroups: (1) antioxidant defense, (2) aggregate clearance, (3) membrane interactions and signal transduction, and (4) glycolysis. The first group of activated proteins/enzymes in antioxidant defense is represented by eight genes. **OSGIN1** codes for oxidative stress-induced growth inhibitor 1, a protein that facilitates the translocation of the p53 protein into the nucleus, thereby contributing to the inhibition of the cell cycle and mobilizing efforts to combat oxidative stress (Brennan et al. 2017). Interestingly, the activation of this gene is quite pronounced. Sulfiredoxin-1 (**SRXN1**) is repairing oxidative damage to cystines by reducing cysteine-sulfinic acid; the activation of SRXN1 has been also reported for diethylmaleate and TBHQ in (Bischoff et al. 2019) and thus, it is an Nrf2 target gene; **SLC7A11 -** a cystine-glutamate antiporter, which allows the entry of one cystine dimer molecule and the exit of one glutamate molecule, and whose inhibition models oxidative stress *in vitro* (glutathione depletion model currently known as ferroptosis); **TXNRD1** - thioredoxin reductase 1, a cytoplasmic selenium-containing enzyme repairing oxidized cysteines in proteins, and supposedly, a target for the ferroptosis inducer RSL3 in addition to GPX4 (Cheff et al. 2023); **GSR –** glutathione reductase - belongs to the same family as **TXNRD1,** but does not contain an additional selenodithiol; **NQO1** codes for an inducible cytosolic NAD(P)H-quinone oxidoreductase 1, which reduces quinones to hydroquinones in a two-electron process, bypassing the radical one-electron-reduced forms of such quinones as ubiquinone. In addition to the apparent detoxification of quinones, the enzyme plays a crucial role in the production of NAD for the enzyme’s poly (ADP-ribose) polymerase (PARP) and sirtuins. Additionally, it interacts with several “initially unstable” proteins, including p53 and the transcriptional activator PGC-1α, thereby preventing their degradation by the 20S proteasome. Currently, the enzyme is considered a cellular redox switch, interacting with several proteins and mRNAs depending on the cell’s redox status (Ross and Siegel 2017). **GCLM** - the regulatory subunit of gamma-glutamylcysteine synthetase, an enzyme that catalyzes the rate-limiting step in the synthesis of glutathione; **HMOX1** – the inducible heme oxygenase 1, a well-known target for Nrf2/Bach1 couple, and usually referred to as protective, since it decomposes free heme, however, recent works demonstrate that it, in fact, accelerates ferroptosis by producing ferrous iron [see (Soni et al. 2024) and references therein]. Interestingly, its induction by peptide is very modest compared to other Nrf2 targets, likely due to existing inhibition from Bach1 for this gene. **LUCAT1** - long noncoding RNA - was recently shown to play a protective role in oxidative stress injury, inflammation, viability, and apoptosis of cardiomyocytes induced by H_2_O_2_ via regulating miR-181a-5p (Xiao et al. 2021).

**Table 1.** Major upregulated genes for cell-permeable fumarate linked Nrf2 peptide.

| Gene | Gene Product | Nrf2-Fumarate Peptide | HPPE |
| --- | --- | --- | --- |
| ***Antioxidant Defense*** |  |  |  |
| OSGIN1 | Oxidative stress induced growth inhibitor 1 | 13.90 | 12.30 |
| SRXN1 | Sulfiredoxin-1 | 7.47 | 3.19 |
| SLC7A11 | Cystine-glutamate antiporter | 6.37 | 11.30 |
| TXNRD1 | Thioredoxin Reductase | 4.54 | 3.34 |
| NQO1 | NAD(P)H-quinone oxidoreductase 1 | 4.25 | 5.61 |
| GCLM | Gamma-glutamyl cysteine synthetase | 3.87 | 5.90 |
| HMOX1 | Heme oxygenase 1 | 2.18 | 64.7 |
| GSR | Glutathione Reductase | 2.14 | 1.69 |
| LUCAT1 | Lung cancer-associated transcript 1 | 3.32 | 14.7 |
| ***Protein aggregate clearance*** |  |  |  |
| SQSTM1 | Ubiquitin-binding protein p62 | 2.51 | 2.41 |
| DNAJB4 | Heat shock protein 40 (homolog) | 2.28 | 2.76 |
| PSMA6 | Proteasome Alpha-subunit type 6 | 2.25 | <1.5 |
| RPS23 | Small ribosome protein 23 | 1.90 | <1.5 |
| GABARAPL1 | GABA type A receptor associated protein like 1 | 1.74 | 1.93 |
| ***Signal transduction and membrane transport*** |  |  |  |
| P2Y6 | Pyrimidinergic receptor P2Y6 | 2.64 | 3.33 |
| PANX2 | Pannexin 2 | 2.44 | 2.50 |
| PALM3 | Paralemmin-3 | 2.30 | 1.75 |
| ABCC3 | ATP binding cassette subfamily C member 3 | 1.73 | 4.93 |
| ABCB6 | ATP binding cassette subfamily B member 6 | 1.74 | 1.98 |
| ***Glycolysis*** |  |  |  |
| ALDOA | Aldolase A | 2.98 | <1.5 |
| ME1 | Malic Enzyme 1 | 2.05 | 2.33 |

| **Feedback regulation** |  |  |  |
| --- | --- | --- | --- |
| BACH1 | Transcriptional repressor of Nrf2 | 1.58 | 2.01 |
| BACH2 | Transcriptional repressor of Nrf2 | 1.59 | 2.04 |

In addition to the above genes, a set of genes encoding proteins responsible for recognizing and clearing aggregated proteins was identified. **SQSTM1 (p62)** is a Keap1 client protein, known as a cellular “garbage collector” that binds to ubiquitinated protein aggregates and delivers them to the autophagy machinery for disposal. Disruption in the p62-autophagy system is a hallmark of neurodegenerative diseases such as Alzheimer’s, Parkinson’s, Huntington’s, and amyotrophic lateral sclerosis (Ma, Attarwala, and Xie 2019). **DNAJB4** encodes a heat shock protein 40 homolog, known to interact with E-cadherin (Simoes-Correia et al. 2014), the mu-opioid receptor OPRM1, and protein SDIM1 (Lei et al. 2011); the downregulation of the latter is associated with Alzheimer’s disease. It likely functions as a chaperone and exhibits anti-apoptotic properties (Shu et al. 2018). **PSMA6** encodes the alpha-6 subunit of the 20S proteasome, whose inhibition leads to proteasome dysfunction, as observed in diabetic nephropathy (Feng et al. 2018), and is a potential target of the Nrf2 transcription factor. **RPS23** codes for small ribosomal protein 23, a hub protein associated with the formation of neurofibrillary tangles in AD (Nguyen, Kim, and Huong Vu 2024). It is a substrate for prolyl hydroxylase OGFOD1 (Xie et al. 2024), and when hydroxylated, it loses its protective function; the OGFOD1 enzyme inhibition with Roxadustat (FG-4592) is, in fact, the main reason for the drug’s protective effect in ischemia-reperfusion (Singleton et al. 2014). **GABARAPL1** is an autophagy-associated protein and, like GABARAP, promotes GABA_A_ receptor clustering (Chen et al. 2000). Low expression of GABARAPL1 decreases sensitivity to ferroptosis in cancer stem cells (Du et al. 2022). The **P2Y6** gene encodes the P2Y_6_ receptor, which appears to mediate Aβ- and tau-induced neuronal and memory loss through the microglial phagocytosis of neurons (Puigdellivol et al. 2021). P2Y6R may have a detrimental or beneficial role in the nervous system in the context of neurological pathologies, such as ischemic stroke, Alzheimer’s disease, Parkinson’s disease, radiation-induced brain injury, and neuropathic pain (Anwar, Pons, and Rivest 2020).

The third group of targets includes signal transduction and membrane interaction players. **PANX2 (Pannexin 2)**, located at the interface between mitochondria and the endoplasmic reticulum (Le Vasseur et al. 2019), is particularly abundant in the brain. A link between Nrf2 activation and PANX2 induction under the action of oltipraz was reported in (Liao et al. 2020), and PANX2 was confirmed as an Nrf2 target gene and one of the risk genes for autism spectrum disorder (Park et al. 2024). **PALM3** codes for parallemmin-3, a phosphoprotein with prenyl and palmitoyl substituents, providing its interaction with the cytoplasmic side of the cell membrane and thus participating in neuronal membrane dynamics (Chen et al. 2011). **PALM3** was never reported as an Nrf2 target gene. **ABCB6** gene codes for an ATP-dependent transporter that catalyzes the transport of a broad spectrum of porphyrins through the plasma membrane and may also function as an ATP-dependent importer of porphyrins from the cytoplasm into the mitochondria and participate in the de novo heme biosynthesis regulation and the coordination of heme and iron homeostasis. ABCB6 knockout mice exhibit an increased auditory brainstem response threshold, resulting in reduced hearing sensitivity, thus pointing to the ABCB6 role in inner ear development and function (Buccheri et al. 1986). **ABCC3** gene codes for an ATP-binding cassette transporter, also known as multidrug resistance-related protein 3, which is involved in the physiological regulation of bile salt enterohepatic circulation and acts as an overflow pump for bile acids as well as glucuronide conjugates of bilirubin and drugs (Patel, Taskar, and Zamek-Gliszczynski 2016). Aldolase A (**ALDOA**) and malic enzyme (**ME1**) are well-known Nrf2 targets.

The transcriptomic analysis of the Nrf2-fumarate peptide suggests its benign character, which is uncommon among chemical activators that work via Keap1 covalent modification. The list of genes for the cell-permeable Nrf2 peptide can be used as a set of “antioxidant pathway only” genes compared to other Nrf2 activators. The genes for Bach1 and Bach2 transcriptional repressors are modestly activated (Table 1). This observation again highlights the feedback regulation in Nrf2 activation through the activated expression of its transcriptional repressor (Jyrkkanen et al. 2011). Therefore, for sustained activation of Nrf2, which is essential for treating chronic diseases, Bach1 inhibition may be just as critical as Nrf2 activation alone.

### 3.6. Comparative transcriptomic analysis reveals that HPPE is both an Nrf2 activator and a Bach1 inhibitor

One of the recently identified HMOX1 activators that exhibit both Nrf2 activation and Bach1 inhibition activity (Ahuja et al. 2021) is HPPE, the best hit among the series of fused benzimidazoles patented by High Point Pharmaceuticals LLC (Attucks et al. 2014). The compound is not an alkylating agent on its own. However, it is active in the Neh2-luc assay and thus directly stabilizes the Nrf2 protein (Ahuja et al. 2021). The mechanism of this effect is currently unknown and is the focus of our research in the laboratory. The recently proposed mechanism of Nrf2 activation based on the HPPE property to act as a zinc ionophore (Freeman and Bollong 2024) is doubtful, as no zinc effect was observed in the Neh2-luc reporter for HPPE-induced activation (Fig. S2). To evaluate whether the transcriptomic landscape of HPPE-treated cells aligns more closely with that of classical Nrf2 activators, such as cell-permeable Nrf2 peptides or Bach1 inhibitors, we employed RNA-sequencing analysis.

A comparative transcriptomic analysis of HPPE action in a neuroblastoma cell line demonstrates that HPPE possesses both activities. The Venn diagrams for HPPE and the two porphyrins (Fig. 8A, B) show quite an intersection with Bach1 inhibitors (71 common activated genes in Fig. 8A, see the list in Table 2) and some intersection with Nrf2-peptide (22 upregulated common genes in Fig. 8C). The list of these genes matches “Nrf2 only” target genes identified with cell-permeable fumarate linked Nrf2 peptide (see Table 1). The K-mean clustering analysis was performed using the empirical number of the clusters by elbow plot (Fig. S3A), which demonstrated that except for a small population of genes in cluster number 3 (Fig 9A dotted box: Fig. 9B). This gene signature in cluster 3 was enriched for canonical Nrf2-Bach1 pathways (Fig S3B). However, the transcriptional landscape of HPPE-treated cells predominantly matched with the transcriptional landscape of zinc and tin-protoporphyrins than those compared to the Nrf2-peptide-treated cells in the top 2000 variable genes (Fig. 9A). Interestingly, cluster 4 with Bach1 inhibition effects (HPPE and protoporphyrin’s) was enriched for neuroactive ligand-receptor interactions (Fig. S3B) indicating a non-canonical role of Bach1 in neuronal physiology independent of Nrf2. These results were further confirmed by employing weighted gene co-expression network analysis (WGCNA) in all observed modules (Fig. S4A-D). Differential gene expression analysis using DEseq2 between HPPE, Nrf2 peptide, zinc and tin-protoporphyrin (FDR q <0.1, Log2FC=2) demonstrated 1577 differentially regulated genes between HPPE and Nrf2-peptide treated cells, whereas 9 differentially expressed genes between HPPE and (Zn & Sn-protoporphyrin) (Fig. 9C), further confirming the similarity of HPPE with Bach1 inhibitors. Overall, the transcriptomic analysis demonstrates that HPPE is a Bach1 inhibitor, very similar to known metal protoporphyrin inhibitors of Bach1, and it also activates the Nrf2 genetic program, like the cell-permeable Nrf2 peptide.

**Fig. 8.**
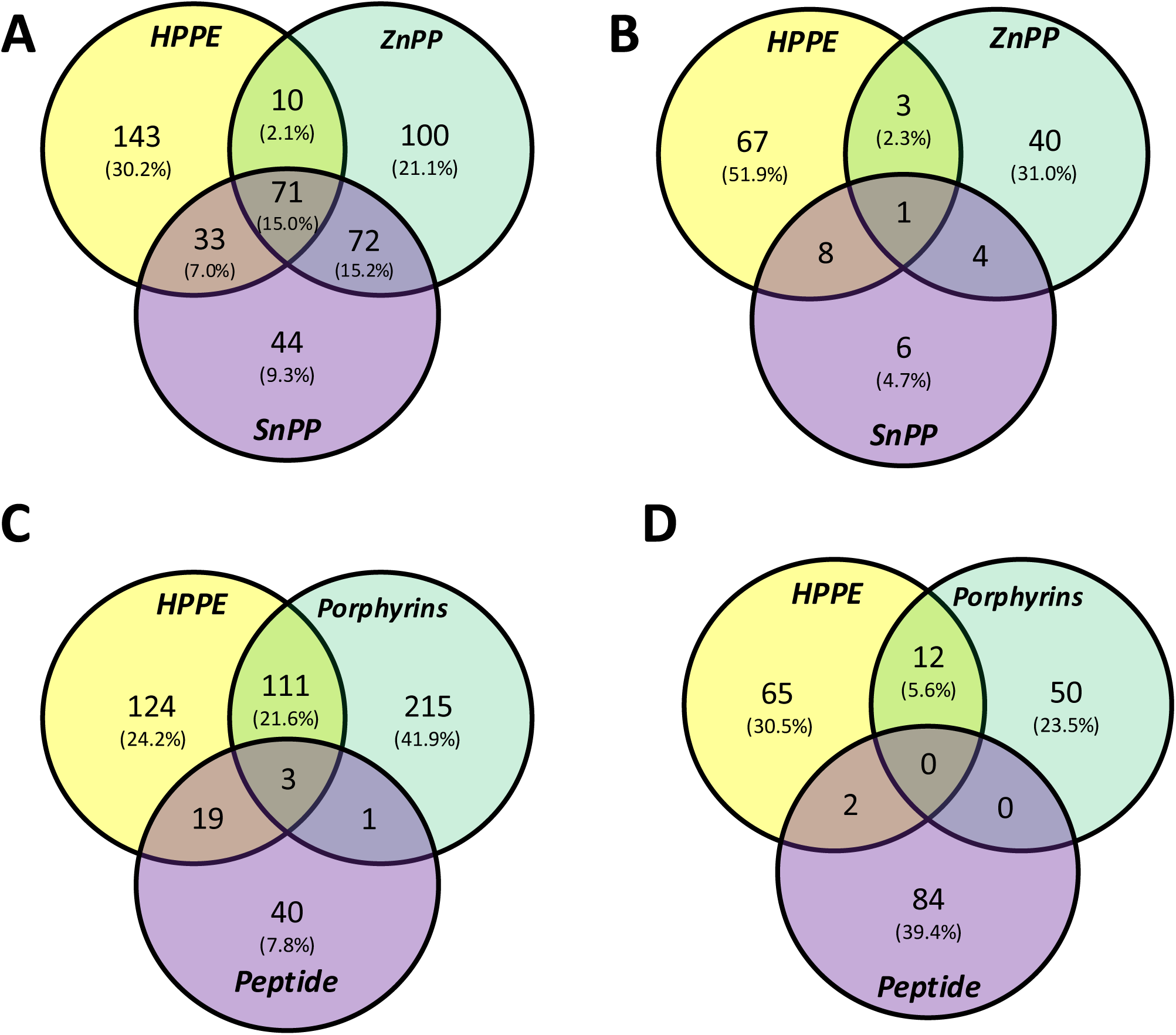
Venn diagrams for upregulated (A, C) and downregulated (B, D) genes for neuroblastoma cell line treated for 5 h with the following compounds: HPPE, Zinc (ZnPP) and tin (SnPP) protoporphyrins at 5 μM each and 100 μM of cell-permeable fumarate-linked Nrf2 peptide.

**Fig. 9.**
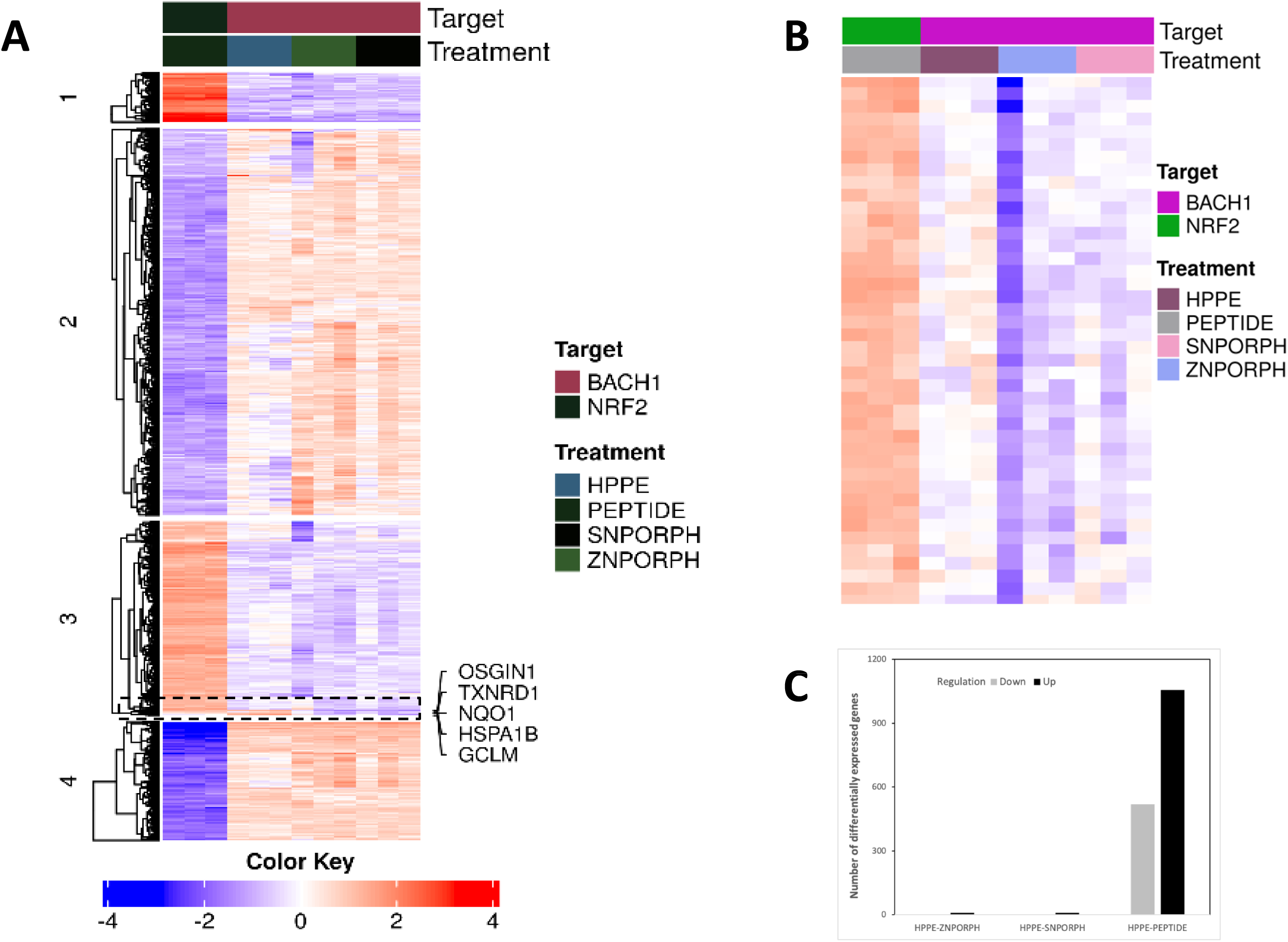
Transcriptomic analysis differentiates gene signatures between Nrf2-peptide and Bach1 inhibitors. (A) K-means clustering of 2000 variable genes. (B) Magnified region from the dotted box in cluster 3 in (A), annotation of the color bar is the same as in (A). (C) Differentially expressed genes (FDR q <0.1, Log2FC=2) between HPPE vs Nrf2 peptide, HPPE vs Zn protoporphyrin, and HPPE vs Sn-protoporphyrin.

**Table 2.**
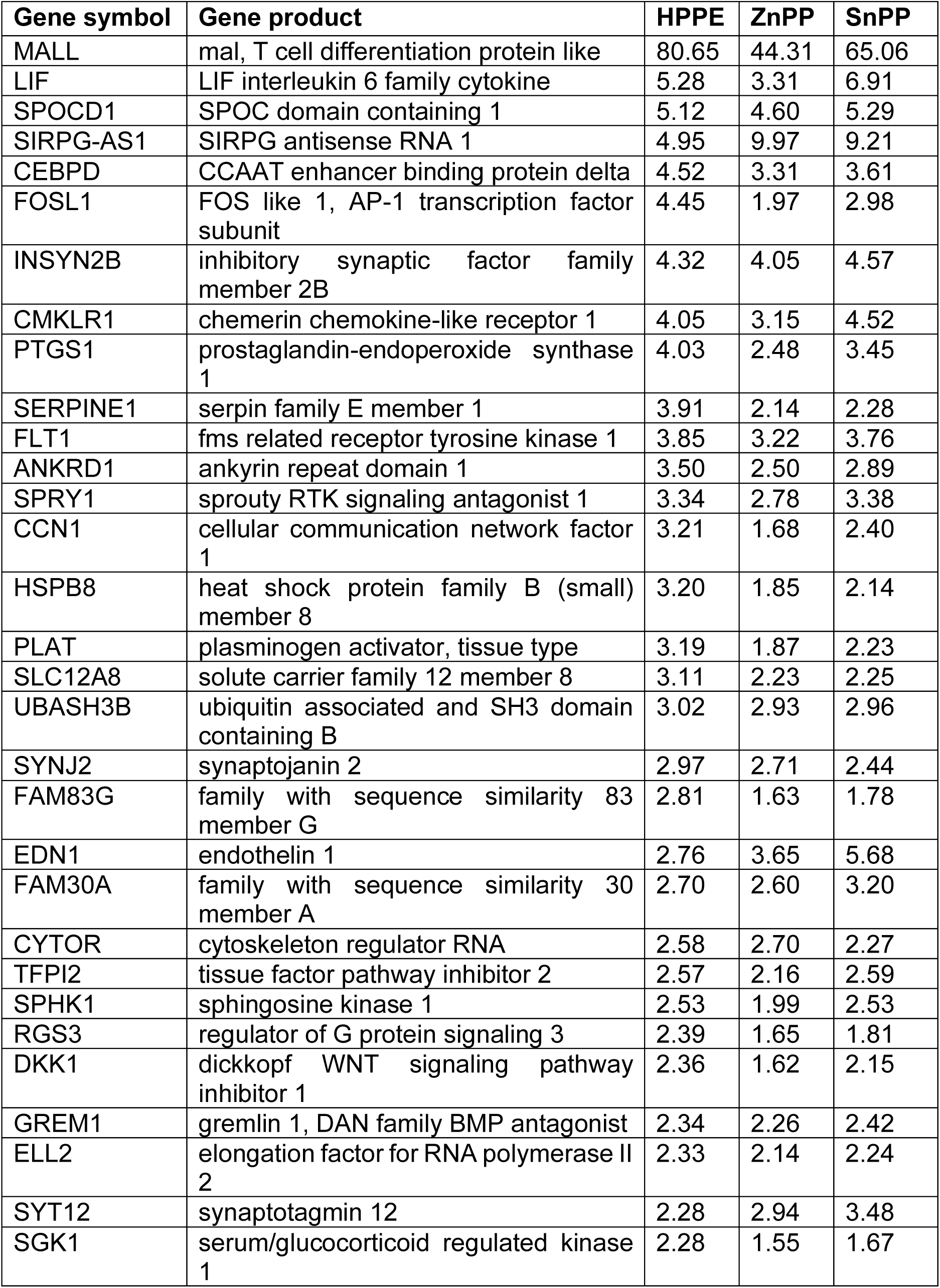
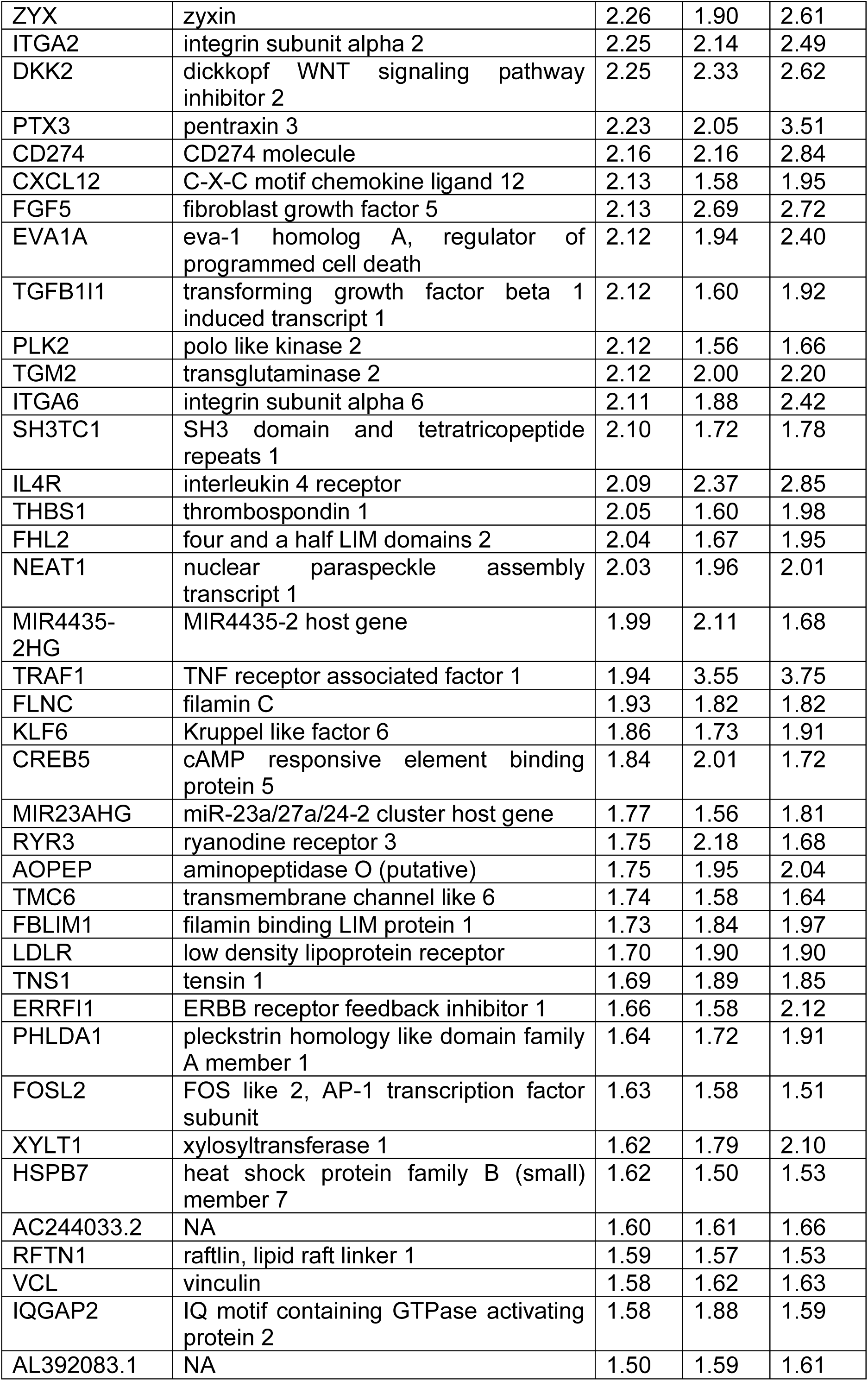
Overlapping upregulated genes for HPPE, zinc, and tin protoporphyrins.

## 4. Conclusions

Aging is associated with a gradual accumulation of lipoxidative and nitrosative byproducts, especially in the central nervous system (Diaz et al. 2024). In addition, aging also leads to fundamental deterioration in homeostatic functions including the regulation of proteostasis, genomic stability, mitochondrial function, telomere maintenance, autophagy, nutrient sensing, cell-cell communication, and cellular senescence (Lopez-Otin et al. 2013). These effects are further amplified by neurodegenerative diseases, where oxidative stress often propagates and sustains the disease progression (Schmidlin et al. 2019).

Nrf2-related genes and oxidative stress regulate these aging-related attritions (Schmidlin et al. 2019) (Zinovkin, Kondratenko, and Zinovkina 2022). Our measurement demonstrates tissue-specific deterioration in the Nrf2 defense during aging (Fig 4C, D, and E). The Nrf2 level in the brainstem, a region critically vulnerable to Parkinson’s disease, was significantly lower in the aged mouse, tipping the Nrf2-Keap1 ratio further in favor of Keap1. The spinal cord also lost Nrf2 expression during aging, while cortical levels of both Nrf2 and Keap1 remained unscathed. Previous results have depicted contradictory effects of aging on the expression of Nrf2 in experimental animals (Zinovkin, Kondratenko, and Zinovkina 2022). In general, brain and spinal cord samples from aging mice showed a decline in Nrf2 activity (Diaz et al. 2024), (Davinelli et al. 2014) except for a few that depicted the opposite (Zhang et al. 2012). Our results also suggest that the relative expression of Nrf2 and Keap1, rather than either protein in isolation, is crucial for understanding physiological relevance. Therefore, despite the aging-related dip in Nrf2 expression in the spinal cord, the relevance of Nrf2 downregulation in the midbrain is more significant. The uneven loss of the Nrf2 activity can elicit region-specific susceptibility to oxidative stress in the brain. Intriguingly, the midbrain, which is most susceptible to the loss of Nrf2 expression, is also the most vulnerable region to neuronal loss in PD, a disease strongly associated with oxidative stress (Dias, Junn, and Mouradian 2013).

The development of Nrf2 activators suitable for treating chronic age-related conditions and diseases necessitates addressing several issues stemming from the mechanistic aspects of Nrf2 stabilization and activation. First, a high intracellular concentration of Keap1 protein sets the limit for the efficiency of reversible displacement activators, as their EC_50_ will always exceed the 1 μM level, independent of their K_D_ value. Second, Keap1 is an adaptor protein for a dozen client proteins for the Cullin III ubiquitin ligase complex. As such, the displacement activator must outcompete endogenous Nrf2 and all other client proteins, which provides an additional offset in EC_50_ values, shifting them to ca. 5-10 μM or above. Third, as we discussed earlier, there may be a significant abundance of other off-target Kelch-domain proteins that could interact with a displacement activator designed to interact with the Kelch domain (Hushpulian et al. 2021). All these factors contribute to the observed offset of biologically effective concentrations of displacement activators by many orders of magnitude compared to their binding constants determined in an FP assay. The redox-sensing nature of Keap1 is a distinct characteristic of this protein. Therefore, a combination of a displacement scaffold with a pro-oxidant or alkylating motif may solve the problem of Keap1-specific targeting and efficient inhibition. This approach has been illustrated using the cell-permeable Nrf2 peptide, which is linked to fumarate through amide bonding to the lysine at the peptide’s C-terminus. The peptide acts as a “pure Nrf2 activator,” upregulating only the Nrf2 genetic program, as confirmed by transcriptomic analysis.

Another problem associated with Nrf2 activation is that such activation has a feedback regulation not only through Keap1, as a well-known gene target of Nrf2, but also through Bach1 and Bach2, transcriptional repressors of Nrf2 and Nrf2 target genes. Therefore, to treat chronic conditions where sustained activation of the antioxidant program is necessary, the ideal Nrf2 activator must exhibit the properties of targeted inhibition of Keap1 and Bach1. The recently developed HMOX1 inducer from the group of benzothiazolyl-amino-benzimidazoles - HPPE – exhibits both activities, e.g., Bach1 inhibition and Nrf2 activation, confirmed by the comparative transcriptomic analysis. The mechanism of Nrf2 activation by HPPE still requires clarification, as the compound lacks alkylation capacity. Nevertheless, the ongoing trials of HPPE and its variants have shown extremely promising results in oxidative stress scenarios (Ahuja et al. 2021).

Dmitry M. Hushpulian - Investigation, Visualization, Data Curation, Conceptualization, Writing – original draft. Navneet Ammal Kaidery – Investigation, Visualization, Conceptualization, Data Curation, Writing – original draft. Priyanka Soni - Investigation, Data curation. Andrey A. Poloznikov – Investigation, Resources, Writing – review & editing. Arpenik A. Zakhariants - Investigation, Data curation. Alexandra V. Razumovskaya - Investigation, Data curation. Maria O. Silkina - Investigation, Data curation. Vladimir I. Tishkov - Writing – review & editing. Eliot H. Kazakov - Investigation, Data curation. Abraham M. Brown - Writing – review & editing, Conceptualization. Irina N. Gaisina – Investigation, Writing – review & editing, Conceptualization. Young-Hoon Ahn - Investigation, Data curation. Sergey V. Kazakov - Writing – review & editing. Nancy Krucher - Writing – review & editing, Resources. Sudarshana M. Sharma - Investigation, Writing – review & editing, Conceptualization, Data curation. Bindu D. Paul- Writing – review & editing, Resources. Irina G. Gazaryan: Writing – original draft, Writing – review & editing, Conceptualization, Methodology, Resources. Sergey V. Nikulin – Investigation, Writing – original draft, Writing – review & editing, Conceptualization, Supervision, Resources. Bobby Thomas- Writing – original draft, Writing – review & editing, Conceptualization, Methodology, Resources, Formal analysis, Supervision, Funding acquisition.

## Data Availability

All data supporting the findings of this study are available within the article and its supplementary information. The RNA-seq data generated in this study are available in the Gene Expression Omnibus (GEO) repository (GSE271364 and GSE287793).

## Declaration of Competing Interest

The authors declare that they have no known competing financial interests or personal relationships that could have appeared to influence the work reported in this study.

## Funding

The work was supported by grants funded by the National Institutes of Health (R01AG077396, R01NS101967, and R01NS133688) and the Department of Defense (HT94252310443) to BT. This work was supported by the Basic Research Program at the National Research University Higher School of Economics to SN, AR, and MS for computer modeling and bioinformatic analysis. SMS is partially supported by the National Institutes of Health grant P01CA203653. BDP is supported by the American Heart Association and Paul Allen Foundation grant 19PABH134580006, R01AG071512, 1R21AG073684, the Solve-ME Foundation, and the Catalyst Award from Johns Hopkins University.

## Acknowledgements

The authors acknowledge Intelligenomica LLC for the genomic analysis support and graphics generation.

## Supplemental materials

**Fig. S1.** Chemical structure of Nrf2 displacement activators with pro-oxidant motifs.

**Fig. S2.** No effect of zinc ion or N-acetylcysteine (NAC) presence at Neh2-luc reporter activation induced by HPPE.

**Fig. S3.** Transcriptomic analysis differentiates gene signatures between Nrf2-peptide and Bach1 inhibitors. (A) Elbow-plot used to determine the number of clusters for K-mean clustering. (B) Enrichment tree of genes represented in clusters 1, 3, and 4 of Fig. 9A using Enrichr.

**Fig. S4.** Weighted gene co-expression network analysis (WGCNA). (A) Visualization of normalized expression at 75 quantile used for WGCNA. (B) Mean connectivity and scale of dependence cutoff used for soft-threshold power for WGCNA. (C) Heatmap visualization of co-expression modules identified through WGCNA analysis of RNA-seq data. (D) Cluster dendrogram of WGCNA with merged colors.

**Table S1.** Docking energies for the versions of cell-permeable Nrf2 peptides studied in the work compared to individual Nrf2 peptides and the TAT-peptide.

**Table S2.** Gene list with Euclidean distance metrics of the 2000 most variable genes shown in Fig. 9A.

## Notes

### Competing Interest Statement

The authors have declared no competing interest.

