## Supplementary figures and images for "FUNCTIONAL ANALYSIS OF BIPARTITE NRF2 ACTIVATORS THAT OVERCOME FEEDBACK REGULATION FOR AGE-RELATED CHRONIC DISEASES"

### Supplementary figures

#### Fig. S1

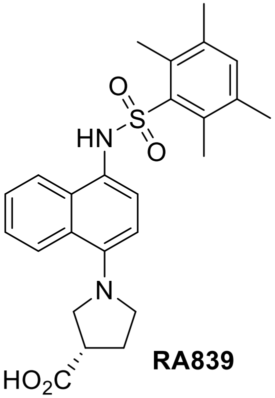

**RA839**

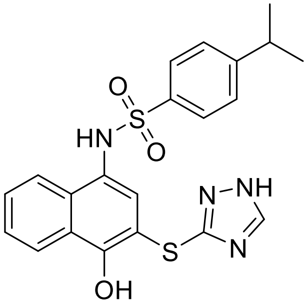

**S47**

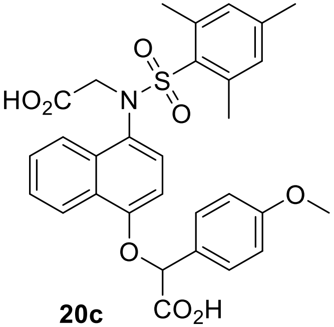

**20c**

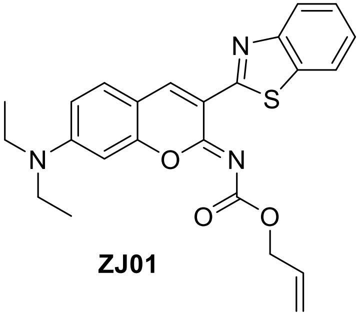

**ZJ01**

**Fig. S2**

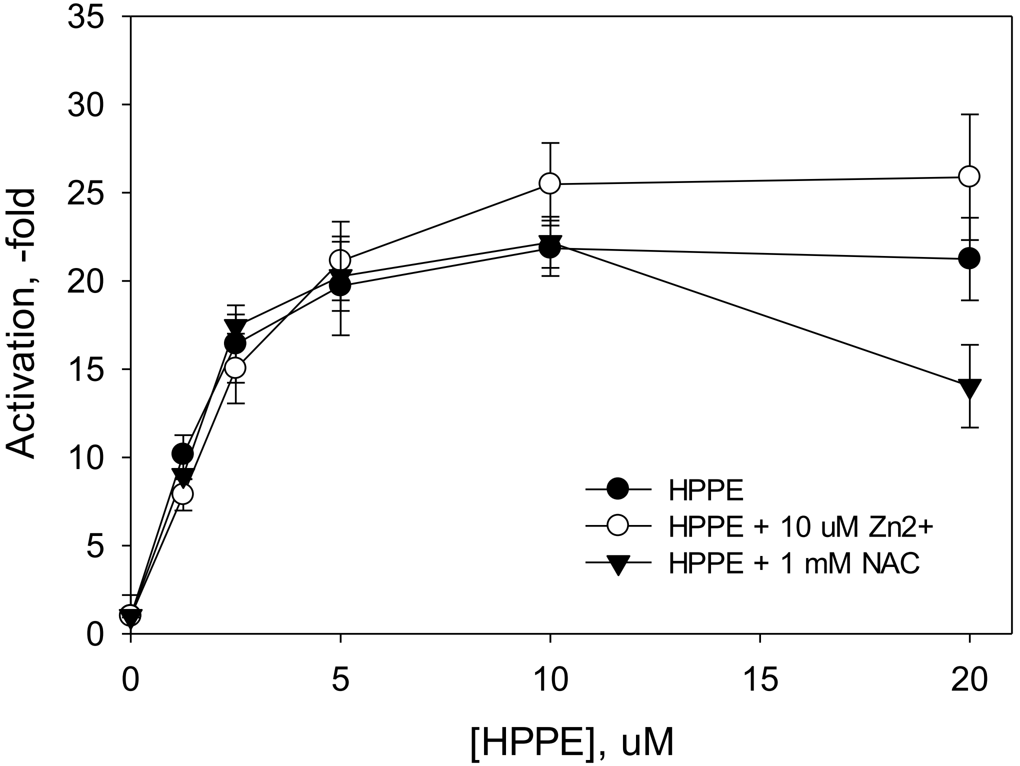


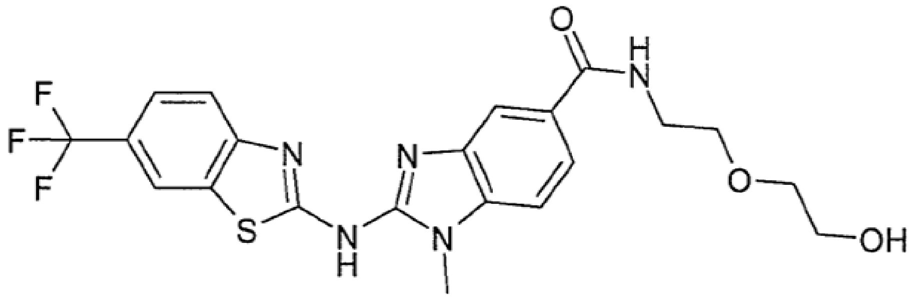

HPPE

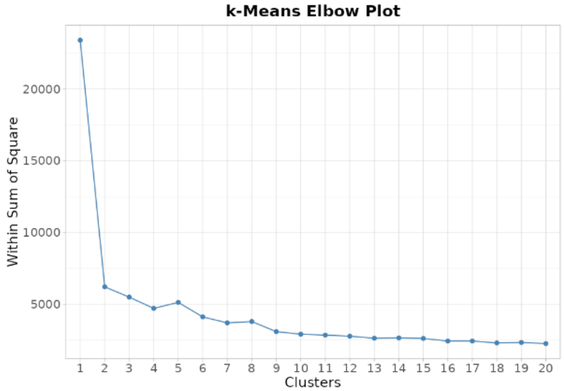

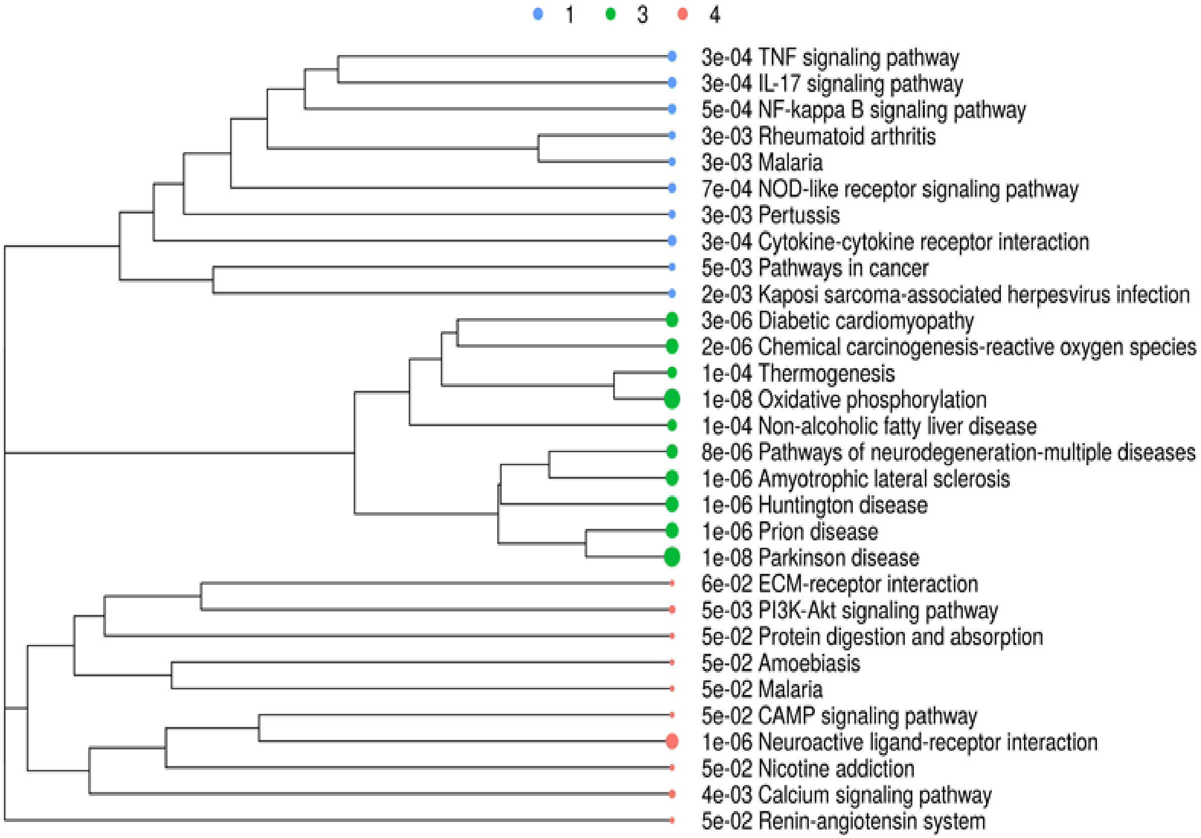

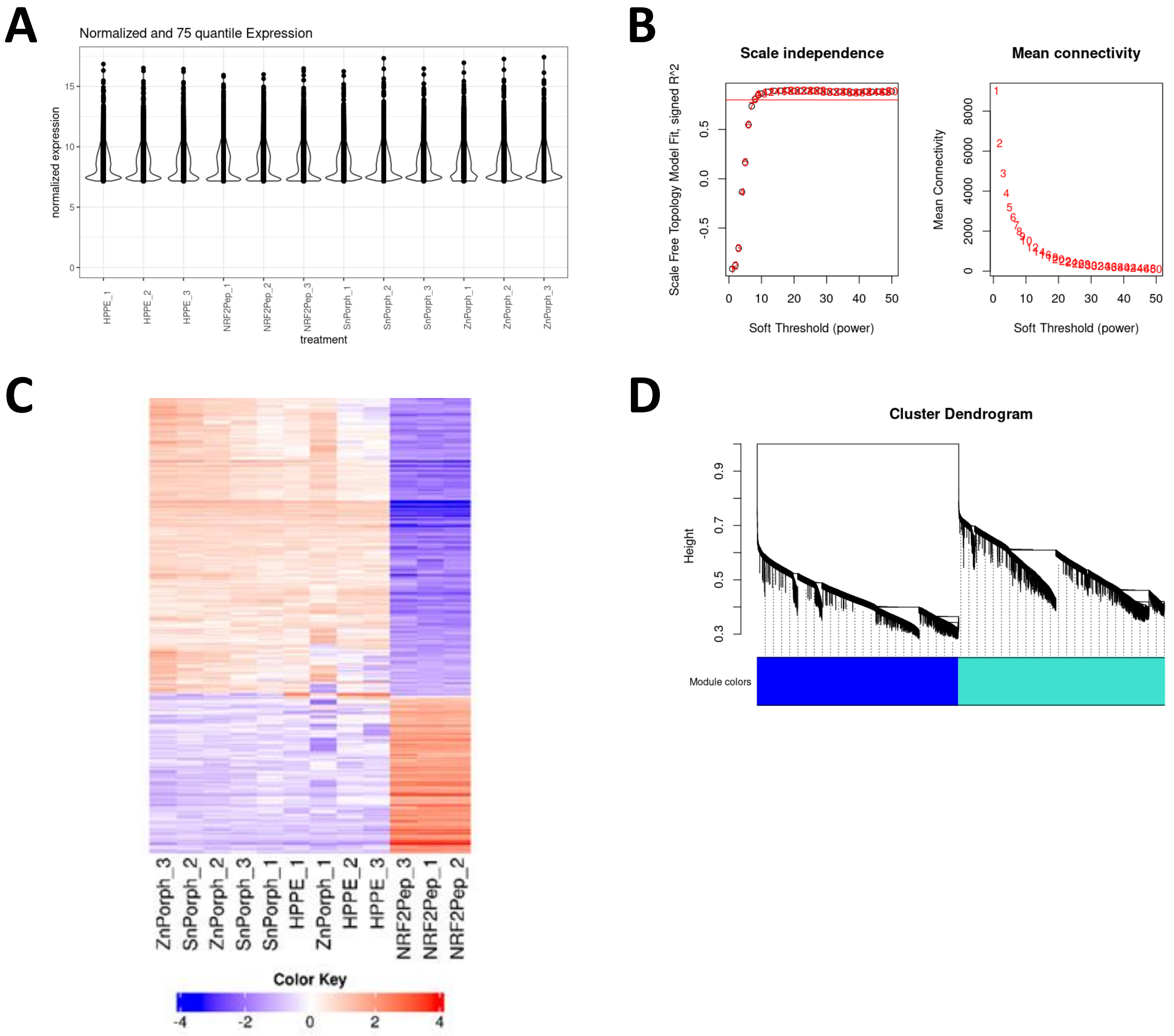
