## Supplementary material for "FUNCTIONAL ANALYSIS OF BIPARTITE NRF2 ACTIVATORS THAT OVERCOME FEEDBACK REGULATION FOR AGE-RELATED CHRONIC DISEASES": Table S1

### Docking benchmark

| Ligand Name | Binding Energy (kcal/mol) | Total Pi Interactions | Total Hydrogen Bonds | Total Salt Bridges | Ligand Contact Surface Area | Receptor Contact Surface Area | Receptor Polar Contact Surface Area | Receptor Nonpolar Contact Surface Area |
|---|---|---|---|---|---|---|---|---|
| TAT ETGE WT Peptide | -1209,9514 | 5 | 26 | 1 | 491,84 | 472,74 | 319,54 | 153,2 |
| Mutant Peptide | -1341,0492 | 6 | 29 | 1 | 534,6 | 512,6 | 327,43 | 185,17 |
| Fumarate-Peptide | -1380,2161 | 7 | 29 | 0 | 597,54 | 565,57 | 366,97 | 198,59 |
| TAT-14 | -1544,5982 | 3 | 10 | 2 | 751,02 | 590,29 | 352,12 | 238,18 |
| ETGE peptide (control) | -1311,2046 | 2 | 13 | 0 | 321,56 | 293,14 | 162,68 | 130,46 |
